# Effects of unconscious tactile stimuli on autonomic nervous activity and afferent signal processing

**DOI:** 10.1101/2024.07.30.605926

**Authors:** Mai Sakuragi, Yuto Tanaka, Kazushi Shinagawa, Koki Tsuji, Satoshi Umeda

**Affiliations:** Department of Psychology, Keio University, 2-15-45 Mita, Minato-ku, Tokyo, 108-8345, Japan; Keio University Global Research Institute, 2-15-45 Mita, Minato-ku, Tokyo, 108-8345, Japan

**Keywords:** Subthreshold Tactile Perception, Heartbeat Evoked Potential, Interoception

## Abstract

Autonomic nervous system (ANS) is a mechanism that regulates our internal environment. In recent years, the interest in how tactile stimuli presented directly to the body affect ANS function and cortical processing in humans has been renewed. However, it is not yet clear how subtle tactile stimuli below the level of consciousness affect human heart rate and cortical processing. To examine this, subthreshold electrical stimuli were presented to the left forearm of 43 participants during an image-viewing task, and electrocardiogram (ECG) and electroencephalogram (EEG) data were collected. The changes in the R-wave interval of the ECG immediately after the subthreshold electrical presentation and heartbeat-evoked potential (HEP), the afferent signal processing of cardiac activity, were measured. The results showed that heart rate decelerated immediately after the presentation of subthreshold electrical stimuli. The HEP during stimulus presentation was amplified for participants with greater heart rate acceleration immediately after this deceleration. The magnitude of these effects depended on the type of the subthreshold tactile stimuli. The results suggest that even with subthreshold stimulation, the changes in autonomic activity associated with orienting response and related afferent signal processing differ depending on the clarity of the tactile stimuli.

## 1. Introduction

The organs in our body constantly work to maintain vital functions, whether we are engaged in some activity or asleep. The autonomic nervous system (ANS) regulates many important physiological systems, including cardiac, respiratory, digestive, vasomotor, and endocrine. Through the regulatory functions of the ANS, organs related to these physiological systems are always spontaneously active; however, stimuli from the external environment also influence them in various ways. The semantic information of external stimuli, such as the emotional expression (e.g., Cacioppo et al., 1985) and evaluation of one’s performance (e.g., Crone et al., 2003, 2004) and physical state (Glass, 2001; Hodossy & Tsakiris, 2020; Lehrer & Gevirtz, 2014; McCraty, 2022; Ring et al., 2015; Shaykevich et al., 2015), have a particularly large influence on ANS activity. Even meaningless external stimuli can alter ANS activity due to changes in the tempo of successive stimuli (Bernardi et al., 2006; Fukumoto et al., 2010; Tanaka et al., 2021; K. Watanabe et al., 2017), attention or anticipation of the next input stimulus (Guerra et al., 2016; Ilves & Surakka, 2012; Lacey & Lacey, 1978, 1980; Poli et al., 2007; Simons et al., 1998), and orienting responses that identify the direction of input of external environmental stimuli (Sokolov, 1963).

The effects of changes in the external environment on ANS activity have been examined mainly using audiovisual stimuli because they are easy to create and control. However, recently, the influence of tactile stimuli on the ANS and interoception, that is, the sensation or information processing of the internal environment (Khalsa et al., 2018), has attracted attention (Crucianelli & Ehrsson, 2023). Tactile stimuli on the skin play a fundamental role in homeostatic regulation of our physiological state by monitoring signals generated inside and outside the body (Burleson & Quigley, 2021; A. D. Craig, 2002, 2003; Crucianelli et al., 2022). Information of special tactile stimuli, such as pain, temperature, and gentle touch, reaches the lamina I or solitary bundle tract nucleus in the spinal cord, which connects to the nucleus of homeostasis and interoception (A. D. Craig, 2003, 2009, 2010). Other tactile information reaches the somatosensory and motor thalamic nucleus (Crucianelli & Ehrsson, 2023). These areas are involved in sensory discrimination, stimulus-orienting responses, emotional responses, and motivation and facilitate ANS activity. In addition, tactile information is transmitted to regions that directly interact with the ANS, such as the brainstem, medulla oblongata, and corpus callosum, contributing to changes in heart rate, blood pressure, and blood flow (Penasso et al., 2023).

Numerous studies have reported that suprathreshold tactile stimuli, such as pain, temperature, and gentle touch, affect ANS activity and related physiological indices, including heart rate, respiration, and blood pressure. These studies have shown that different intensities of tactile stimuli elicit changes in the ANS activity in different directions. For example, the feeling of pain due to relatively strong electrical stimuli often produces changes indicative of arousal or sympathetic activation (As reviews, Forte et al., 2022; Koenig et al., 2014). In contrast, the detection of a weak electrical stimulation near the threshold decelerates the heart rate immediately (Grund et al., 2022; Motyka et al., 2019). This heart rate deceleration has been interpreted as being related to parasympathetic activity in the orienting response to environmental changes (Sokolov, 1963). Thus, different physical intensities of stimuli, even of the same type, may elicit responses with different psychophysiological bases.

To our knowledge, few studies have investigated the changes in ANS activity when the intensity of the tactile stimuli is weakened below an individual’s threshold. Previously, we presented a subthreshold vibration stimulus to a participant’s right forearm to change the pulse rate immediately after the presentation (Sakuragi et al., 2023). We found that the participant’s pulse rate fluctuated more significantly when the subthreshold vibration was presented than when no stimuli were presented. However, we could not examine whether the sympathetic or parasympathetic activity was predominant because of the large individual differences in the direction of the pulse rate change. Considering only the closeness of the stimulus intensity, the presentation of subthreshold tactile stimuli is as likely to elicit orienting responses as near-threshold tactile stimuli (Grund et al., 2022; Motyka et al., 2019). However, in these studies, participants knew that tactile stimuli would be presented and allocated attentional resources for stimulus detection. Such awareness and anticipatory cognitive processes can affect autonomic responses. By employing subthreshold tactile stimuli, we can directly examine the effects of tactile stimuli on physiological responses by eliminating conscious perception and prior anticipation of tactile stimuli.

The heartbeat-evoked potential (HEP) is an index that reflects the cortical or subcortical processing of the afferent information of body signals, including the ANS. It is the event-related potential (ERP) evoked by the R-wave of the electrocardiogram (ECG) (Park & Blanke, 2019). HEP amplitudes in the right frontal, right central, and midline regions decrease with the presentation of cold painful tactile stimuli (Shao et al., 2011). However, it is not clear whether the conscious perception of the tactile stimulus or the change in ANS activity associated with pain perception alters HEP. In addition, as HEP can be modulated by emotion (Couto et al., 2015; MacKinnon et al., 2013), it may be influenced by emotional reactions, such as fear and aversion, associated with pain. By examining changes in HEP following the presentation of subthreshold tactile stimuli, we may be able to directly examine the effects of changes in ANS activity on HEP, eliminating the influence of conscious perception of tactile stimuli and accompanying emotional responses.

Therefore, the purpose of the present study was to investigate whether subthreshold tactile stimuli produce changes in ANS activity and the processing of physiological changes by the central nervous system. We used electrical stimuli as the subthreshold tactile stimuli as their physical intensity can be easily fine-tuned. As an index of ANS activity, changes in heart rate were recorded using ECG, which is a simple and noninvasive measurement method. In addition, HEP was used to reflect the cortical or subcortical process of change in ANS activity. This study also exploratorily examined how the intensity and texture of the subthreshold tactile stimuli affect subsequent heart rate changes. An increase in the intensity of suprathreshold electrical stimuli increase the degree of heart rate acceleration and cutaneous vasoconstriction (K. D. Craig & Prkachin, 1978; Kemppainen et al., 1994, 2001; Vassend & Knardahl, 2005). Thus, we assumed that, even for subthreshold tactile stimuli, a higher physical intensity would produce a greater degree of heart rate change. To the best of our knowledge, no previous studies have directly compared the effects of different textures of electrical stimuli on heart rate change using suprathreshold electrical stimuli. Most studies have reported the use of square waves (Al et al., 2021; Grund et al., 2022; Motyka et al., 2019) or pulses similar to square waves with electrical waveforms that rapidly switch between high and low voltages (Kemppainen et al., 1994; Piovesan et al., 2019; Vassend & Knardahl, 2005). Previously, we have shown that subthreshold vibration stimuli of continuously varying intensity over a fixed period can alter the pulse rate; however, the effect of discrete subthreshold stimuli such as square waves is not yet clear. To explore these two factors, subthreshold electrical stimuli consisted of two intensity levels (50% / 90% threshold) and two stimulus types (sine/square waves).

The hypotheses were as follows: (1) subthreshold electrical stimuli would change the heartbeat immediately after presentation; (2) the heart rate change effect would be greater in the 90% threshold condition than in the 50% threshold condition; and (3) the greater the degree of change in heart rate due to the subthreshold electrical stimuli, the greater the amplitude of HEP.

## 2. Methods

### 2.1 Participants

Forty-three undergraduate and graduate students (30 women and 13 men; *M*_*age*_ = 21, *SD*_*age*_ = 1.39) were recruited. The sample size was determined using G*Power 3.1.9.6 (Faul et al., 2007). We performed power analyses to ensure adequate statistical power for the following analyses: 1) one sample *t*-test between the RR interval immediately after the electrical stimulus presentation and 0 and 2) two-way repeated measures analysis of variance (ANOVA) between the types of electrical stimuli (2) × electrical stimulus intensity (2). Considering the expected effect sizes 0.5 and 0.25, respectively, alpha level of 0.05, and power of 0.80, the required sample sizes were 34 and 36, respectively. Adequate data were obtained to allow for the exclusion owing to measurement noise and outliers. To the best of our knowledge, no similar study has reported an effect size; therefore, we assumed a medium effect size as mentioned by Cohen (1988). All the participants had normal or corrected-to-normal vision. This study was approved by the Keio University Research Ethics Committee (no. 220190100) and was conducted in accordance with the Declaration of Helsinki. All participants provided written informed consent before participation. If they had known in advance that subthreshold electrical stimuli would be presented, they might have paid more attention than necessary to the stimulus presentation device and body part to which the stimuli would be presented. In this case, we might not have been able to accurately examine the effects of subthreshold electrical stimulus presentation on heart rate. Thus, we did not explain the presentation of subthreshold electrical stimuli during the image-viewing task in the pre-experimental briefing (see Section 2.3.3). After the experiment, we disclosed the presentation of the subthreshold electrical stimulus during the task, its purpose, and the safety measures taken, and obtained written informed consent again for the use of the data collected through the experiment.

### 2.2 Apparatus

We used Xpod USB pulse oximeter (Nonin 3012LP) and soft-clip cuff (NONIN, 8000SL) placed on the tip of the left index finger of the participants to measure their pulse rates in the heartbeat counting task (see Section 2.3.1).

EEG and ECG activities were recorded during the image-viewing task (see Section 2.3.3). EEG activity was recorded using NetStation 4.5.1, with a 64-channel HydroCel Geodesic Sensor Net at a sampling rate of 500 Hz. Cz was used as a reference during the recording (Yao et al., 2019). ECG signals were measured using Polygraph Input Box (EGI). Ag/AgCl electrodes were attached to the backs of the right wrists and left ankles of the participants. The image stimuli were presented using Presentation software (Neurobehavioral Systems, Inc.), and the subthreshold electrical stimuli, using STM100C (amplifier for adjusting stimulus intensity), STMISOC (constant electrical/constant voltage output device), and EL350 (electrode for electrical presentation) (all BIOPAC). The participants placed their left hand on a cushion (Seria Co., Ltd., Japan), 17 cm wide and 37 cm long, with a polyester surface cover and polypropylene filling. They also wore an electrode for subthreshold electrical presentation inside the left forearm. Stim Tracker (Cedrus) was used to track the timing of the presentation of the image and electrical stimuli, as well as the EEG and ECG data simultaneously. The arrangement of these devices is illustrated in Fig. 1.

**Fig. 1.**
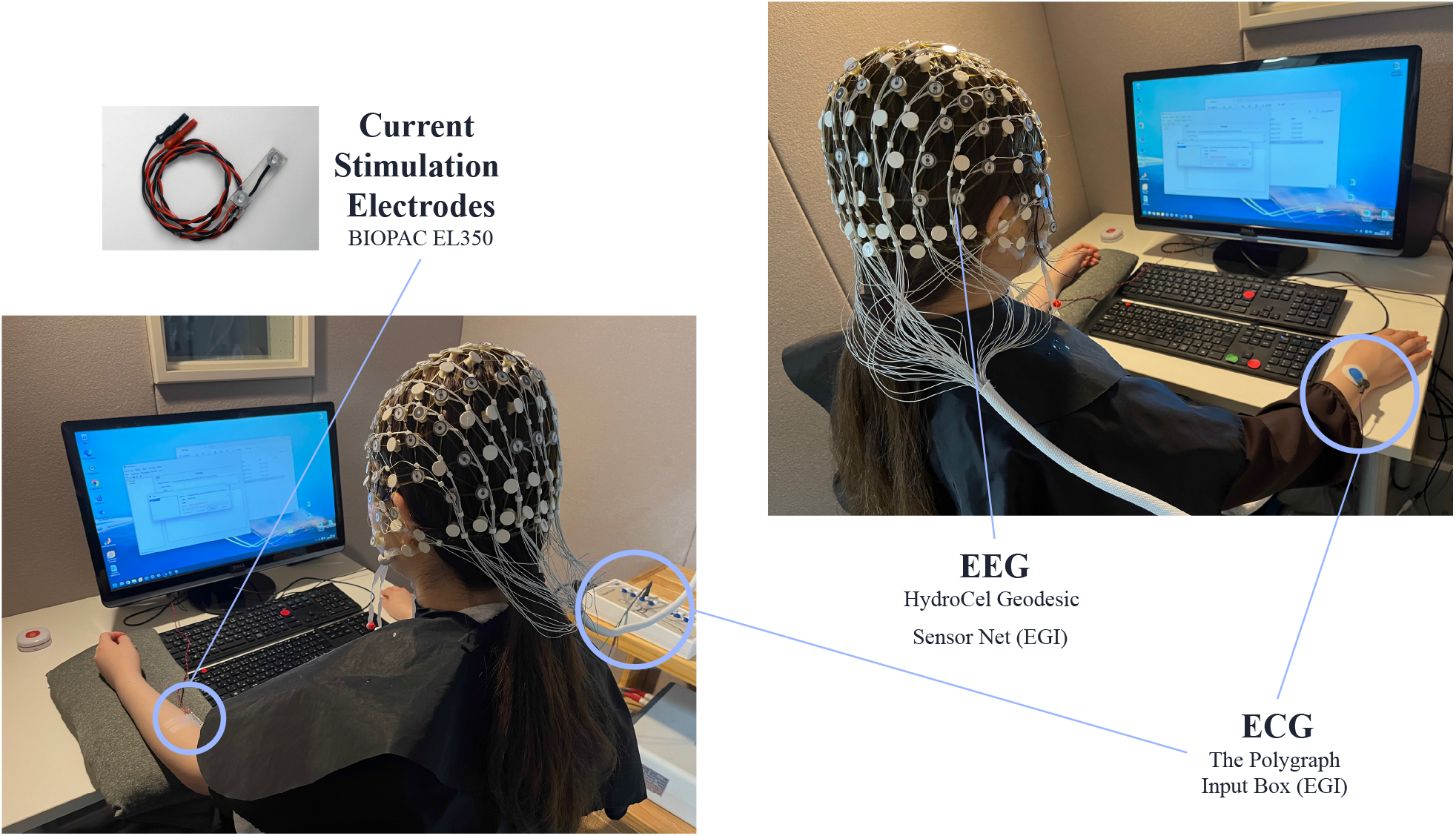
Arrangement of EEG, ECG, and electrodes for subthreshold electrical stimulus presentation. During the image-viewing task, the participants wore an EEG cap and attached ECG electrodes to their right wrists and left ankles. The left arm was placed on a cushion and an electrode was attached to the medial part of the left forearm for subthreshold electrical stimulus presentation. EEG, electroencephalogram; ECG, electrocardiogram.

### 2.3 Procedure

#### 2.3.1 The Heartbeat Counting Task

We used the heartbeat counting task (HCT) to measure the participants’ interoceptive accuracy. This is the objective exactness in detecting internal bodily sensations among the subcategories of interoception (Garfinkel & Critchley, 2013). Cardioception (Legrand et al., 2022), a software that measures cardiac interoception in Psychopy, was used in the experiment. Participants sat in a chair, wore headphones, and silently counted the number of heartbeats between the Start and Stop sounds. They performed trials of 25, 30, 35, 40, 45, and 50 s in a random order. After each trial, they reported the number of heartbeats they had counted and rated their confidence in their performance on a scale of one to seven. They were instructed not to count their heart rate by touching themselves, pressing their bodies hard against a backrest or desk, holding their breath, or relying on pulsation associated with external pressure from headphones. In addition, referring to an adapted version of the teaching of the HCT (Desmedt et al., 2018), participants were asked not to guess and count heartbeats that they did not feel.

#### 2.3.2 Measurement Perception Threshold of Electrical Stimuli

To determine the intensity of the electrical stimuli presented during the main task on an individual basis, the perception threshold of the electrical stimuli was measured for each participant. The psychophysical staircase method was used for the threshold measurement (Fig. 2). The electrical stimulator used could output both sine and square waves; however, we used square waves to measure the stimulus threshold because sine waves are inappropriate for precisely defining the perceptual threshold, as the stimulus intensity varies periodically around a set value. The participants placed an electrode inside their left forearms. When they experienced electrical stimuli, they responded by pressing the F1 key on the keyboard. They performed five trials at one-second intervals for each intensity of the electrical stimulus (one block). A positive response was defined as detecting a stimulus at least once and a negative response, as detecting none. We started with a voltage of 5 V, which was reliably perceived by all the participants and decreased it by 1 V until the first negative response. When a negative response occurred, the voltage for the block was recorded and increased by 0.25 V. When a positive response was obtained in two consecutive blocks, the voltage was lowered by 0.25 V. This procedure was terminated after the voltage was recorded for four negative responses. The average voltage of the four negative responses was used as the perception threshold for the electrical stimuli for each participant. The mean voltage at the participants’ perceptual threshold was 1.855 V, with a standard deviation (SD) of 0.592. Thus, the mean of the voltages presented in the 50% and 90% subthreshold conditions were 0.927 V (upper limit: 1.625 V; lower limit: 0.500 V) and 1.669 V (upper limit: 2.925 V; lower limit: 0.900 V), respectively.

**Fig. 2.**
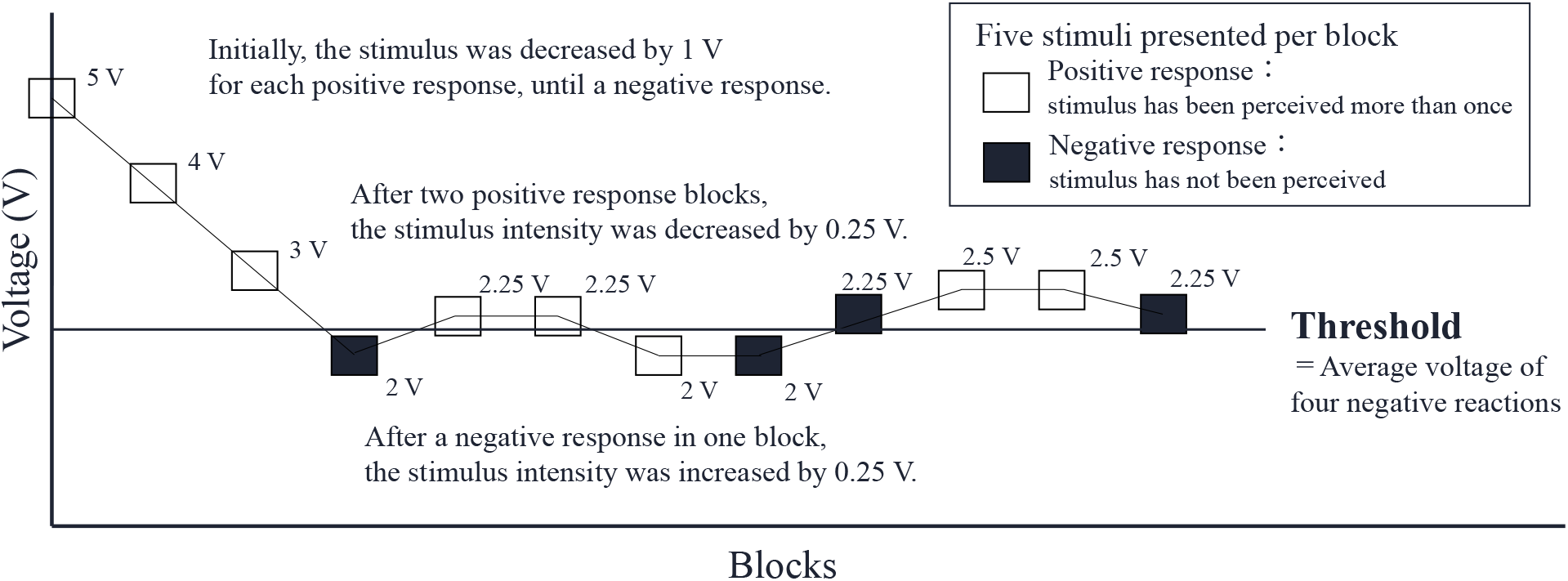
Flow of measuring the participants’ perception threshold of electrical stimuli using the staircase method. The participants responded by pressing the F1 key on the keyboard when they felt an electrical stimulus from an electrode attached to their left forearm. Of the five stimuli of different intensity, a positive response was considered when the stimulus was detected at least once and a negative response, if no electrical stimulus was detected. The stimulus voltage was decreased by 0.25 V if two consecutive positive responses were detected and increased by 0.25 V if even one negative response was detected. The threshold measurements were terminated after four negative responses, and the average voltage of these four negative responses was used as the participant’s threshold value.

#### 2.3.3 Image-Viewing Task (ECG and EEG measurements during subliminal electrical stimulus presentation)

After the perception threshold was measured, the participants wore EEG caps, ECG electrodes, and electrodes for subthreshold electrical stimulus presentation (see Fig. 1). They were instructed to work on a task to measure physiological responses as they viewed images projected on a screen. The experimenter told them that the electrical stimulator on the left arm was a device to measure electrical activity in the skin; however, subthreshold electrical stimuli were presented from this electrode during the task. The first block was the baseline condition, in which no electrical stimulus was presented; only the ECG and EEG signals were measured while viewing the image stimulus. In the following four blocks, subthreshold electrical stimuli of different intensities or types were presented. The four electrical stimuli conditions were prepared by combining two types (sine and square waves) and two intensities (50% and 90% of each participant’s electrical perception threshold). The frequency of the electrical stimuli was set at 140 Hz, following a previous study in which vibration stimuli were presented to the forearm to induce changes in the heart rate (Barralon et al., 2008). In all conditions, the duration of the electrical stimuli was 500 ms and the inter-stimulus interval was 3 s. One image stimulus was presented every 12 s. We used 40 images from the Open Affective Standardized Image Set (OASIS; (Kurdi et al., 2017)) with arousal between 3.0 and 4.0 and valence between 4.0 and 6.5, that is, low arousal and positive emotional valence. Images of human figures and food that are likely to alter the physiological state of humans were excluded.

#### 2.3.4 Image Evaluation Task

After all the tasks were completed, the participants were asked to rate the subjective arousal and valence of the images in the image-viewing task. They first rated the arousal of all stimuli on a 5-point scale from 1 (most calm) to 5 (most intense), followed by valence on a 5-point scale from 1 (most negative) to 5 (most positive), by focusing on the change in their bodily state as a result of viewing the stimuli, rather than on the meaning of the image stimuli. The order in which the images were presented was randomized across the participants.

#### 2.3.5 Questionnaire Survey

The questionnaires used in this study were Somatosensory Amplification Scale (SSAS; Barsky et al., 1990), Body Perception Questionnaire (BPQ-BA; Kobayashi et al., 2021), Multidimensional Interoceptive Sensory Assessment (Japanese Version of MAIA; Mehling et al., 2012), and the Blushing Propensity Scales (BPS; Leary & Meadows, 1991). The scales examined the tendency to perceive bodily sensations as intense, harmful, or disturbing; sensitivity to various sensations and states of one’s body; metacognition of interoception; and tendency to blush in embarrassing situations in daily life, respectively. The SSAS and BPS data were not analyzed or discussed in this study because they were obtained for other research projects. In addition, the participants’ sex and age on the day of the experiment and whether they perceived the subthreshold stimuli during the task on their consciousness were assessed. None of the participants perceived the subthreshold presentation of the electrical stimuli during the experiment.

### 2.4 Pre-processing

#### 2.4.1 Heartbeat Counting Task

Cardioception, which we used for HCT, analyzes the raw pulse wave signal in real-time using a systolic peak detection algorithm and calculates the heart rate as the average of pulse intervals. Using the actual heart rates recorded by this software (Counts) and those reported by the participants (Reported), we calculated the HCT scores per trial for each participant as 1 - |Counts-Reported| / Counts. The mean score of the six trials for each participant was calculated as an index of cardiac interoceptive accuracy. The confidence in each trial was calculated by averaging the values of the six trials, as in the case of the score, and was used as the person’s confidence in cardiac interoception.

#### 2.4.2 EEG and ECG

Offline EEG and ECG data were processed using EEGLAB (version 2023.1, University of San Diego, San Diego, CA, USA). The R-waves of the ECG data were detected using ecglab slow, which is implemented in HEPLAB, a tool in EEGLAB (Perakakis, 2019). Continuous EEG data were down-sampled to 250 Hz and referenced to the common average (Yao et al., 2019). The data were bandpass-filtered between 0.5 and 40 Hz. The noisy EEG channels were interpolated using the pop_interp function in EEGLAB. This function interpolates electrode data containing a large amount of noise using neighboring channel data. In this study, spherical spline interpolation was used, following a commonly used method (Miyakoshi, 2023). The HEP was extracted from 500 to 1500 ms, and epochs with electrical noise were visually removed. Epochs containing noise due to electrical stimuli were not excluded based on the EEG voltage because the degree of noise was different between the participants, even for the same stimulus type or intensity. After visual epoch rejection, the average number of remaining epochs for each stimulus condition was as follows: sine_50, 219; sine_90, 173; square_50, 202; and square_90, 182. We conducted independent component analysis (ICA) using SASICA for artifact removal (Chaumon et al. 2015). To exclude artifacts from the EEG data, ICLabel (Pion-Tonachini et al., 2019), an EEGLAB plugin that automatically determines artifacts, was used. ICA was excluded if it was determined to be muscle, eye, or channel noise with a probability of more than 70% or other artifacts with a probability of more than 80% on ICLabel. To exclude the influence of the ECG cardiac field artifact on the HEP in later model analyses, the mean value of the voltage of the ECG electrode in each stimulus-type condition (base/sine/square) was calculated and included in the model as a parameter (see Section 2.7). Statistical analyses were conducted using R (version 4.3.1). Because of the noise in the EEG data across all epochs for two participants, we excluded their data from the analyses using the EEG data.

#### 2.4.3 Arousal and Valence of Image Stimuli

To examine whether any of the 40 image stimuli employed in the image-viewing task differed significantly from the others in terms of the participants’ subjective ratings of arousal and valence, the distribution of the participants’ ratings for each image was calculated (Appendix Fig. A.1 and A.2). The solid vertical line represents the mean of the ratings for all images and the wavy vertical lines represent the overall mean ± 1SD and the mean ± 2SD. Because the median values of arousal and valence did not exceed the overall mean ± 1SD for any image, it was assumed that no image had a specific effect on the participants’ physical or arousal state during the task. Therefore, experimental data were not excluded from the analysis based on the image stimuli presented during the task.

#### 2.4.4 The Scores of Questionnaires

From the questionnaires, we calculated the mean scores for all BPQ and MAIA items, as well as the MAIA sub-items of noticing ((Q1+Q2+Q3+Q4)/4), attention regulation ((Q11+ Q12+ Q13+ Q14+ Q15+ Q16+ Q17)/7), and body listening ((Q27+ Q28+ Q29)/3) (Mehling et al., 2012).

### 2.5 Group-level ECG Analysis

#### 2.5.1 RR Interval

First, the time interval between R-waves (RR interval) was used to examine the effect of electrical stimuli on the heart rate. The RR interval immediately before the electrical stimulus presentation was designated as RR0 and was considered as an index of normal bodily response before the stimuli presentation. The first and second RR intervals after stimulus presentation were designated as RR1 and RR2, respectively (Fig. 3). For the base condition, the time points of RR0, RR1, and RR2 were set as if the stimulus was presented once every 3 s for 500 ms immediately after the start of the experiment. The average differences between RR1 or RR2 and RR0 (RR1-RR0 and RR2-RR0, respectively) were calculated for the base and four stimulus conditions (hereafter sine_50, sine_90, square_50, and square_90). Because the stimulus interval was 3 s, some participants were presented with the next stimulus before RR3, and the amount of data was not as stable as for RR1 and RR2. Therefore, we did not use time after RR3 as an index in this study. The mean and SD for all participants were calculated for each stimulus condition (base, sine_50, sine_90, square_50, square_90) and time point (RR1-RR0 and RR2-RR0). The participants whose RR difference values exceeded the overall mean ± 3 SD at each time point for each stimulus type were excluded from the analysis. Consequently, we excluded the data of one participant in the sine_50 and square_50 conditions for RR1-RR0. First, to examine the direct effects immediately after the electrical stimulus presentation, one sample *t*-test was performed on the difference between 0 for RR1-RR0 and RR2-RR0 for each condition. The Benjamini-Hochberg (BH) method was used to control for false-positive rates due to multiple comparisons. Moreover, to examine whether the type and intensity of electrical stimuli had different effects on changes in the RR interval, we performed two stimulus intensities × two stimulus types two-way repeated measures ANOVA on the means of RR1-RR0 and RR2-RR0. As the results showed that there were no significant differences in the main effects of stimulus intensity on cardiac activity, stimulus intensity was not used as an independent variable in the following analysis, and physiological data for each participant were calculated for each stimulus type, averaging the values for the two intensities for each participant.

**Fig. 3.**
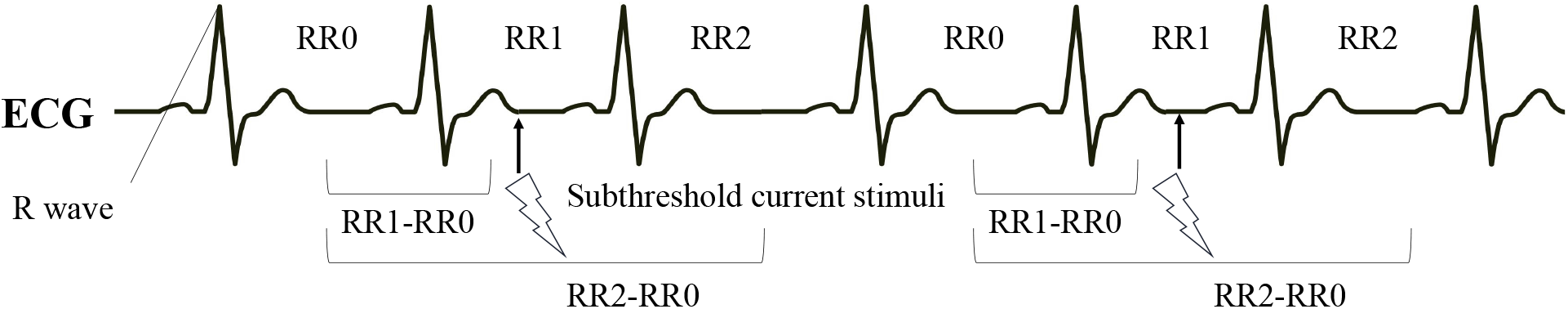
Relationship between the timing of subthreshold electrical stimulus presentation and R-waves of the ECG. The RR interval (time interval between R-waves) immediately before stimulus presentation was RR0, and the two RR intervals immediately after that were RR1 and RR2, respectively. For each participant, we used RR0 as the RR interval without stimulus presentation and the difference between the two RR intervals, RR1-RR0 and RR2-RR0, as a measure of the changes in heart rate immediately after stimulus presentation. RR0, RR interval (time interval between R waves of ECG) immediately before the presentation of subthreshold current stimuli; RR1, RR interval including the time point of current stimulus presentation; RR2, interval immediately after current stimulus presentation; RR1-RR0, the value of the difference between RR1 and RR0; RR2-RR0, the value of the difference between RR2 and RR0.

#### 2.5.2 Correlation of Heart Rate Change Effect with Other Indicators

To examine the factors associated with individual differences in the effects of electrical stimuli on cardiac activity, we evaluated the correlation between the RR1-RR0 and RR2-RR1 values in the sine and square conditions and threshold of the electrical stimuli, score and confidence in the HCT, scores of noticing, attention regulation, and body listening in MAIA, and BPQ scores. The BH method was used for all factor combinations to control for the false-positive rate owing to multiple comparisons. No statistically significant correlation was found (Appendix Fig. A.3-A.9).

### 2.6 Right Frontal HEP Amplitude by Stimulus Type

Using all EEG electrode data, HEP voltage values were averaged every 100 ms from R-wave -200 to +800 ms for the baseline block and each of the two types of electrical stimulation blocks to create a topographic map reflecting the magnitude of the EEG electrode voltage. The topography maps showed that the voltage was negatively amplified in the right frontal region from 200 to 300 ms after the R-wave in all the blocks (Fig. 4A). Therefore, we calculated the HEP using E1, E2, E5, E59, and E60 in the right frontal region, where the electrode is indicated by a black circle in the topographic map in Fig. 4A. HEP was calculated by averaging the amplitude of the EEG voltage from 200 to 300 ms after the R-wave for each block of the base, sine, and square conditions (Fig. 4B). Noise was generated in the EEG immediately after the presentation of the subthreshold current stimuli and its magnitude and length differed between participants and stimulus types. Therefore, when HEPs were additionally averaged for each time point, such as RR1 and RR2, the number of epochs available differed significantly between participants and time points. Therefore, HEP was calculated by summing and averaging all epochs in each block for each participant, reflecting the status of afferent signal processing of the cardiac activity in each block.

**Fig. 4.**
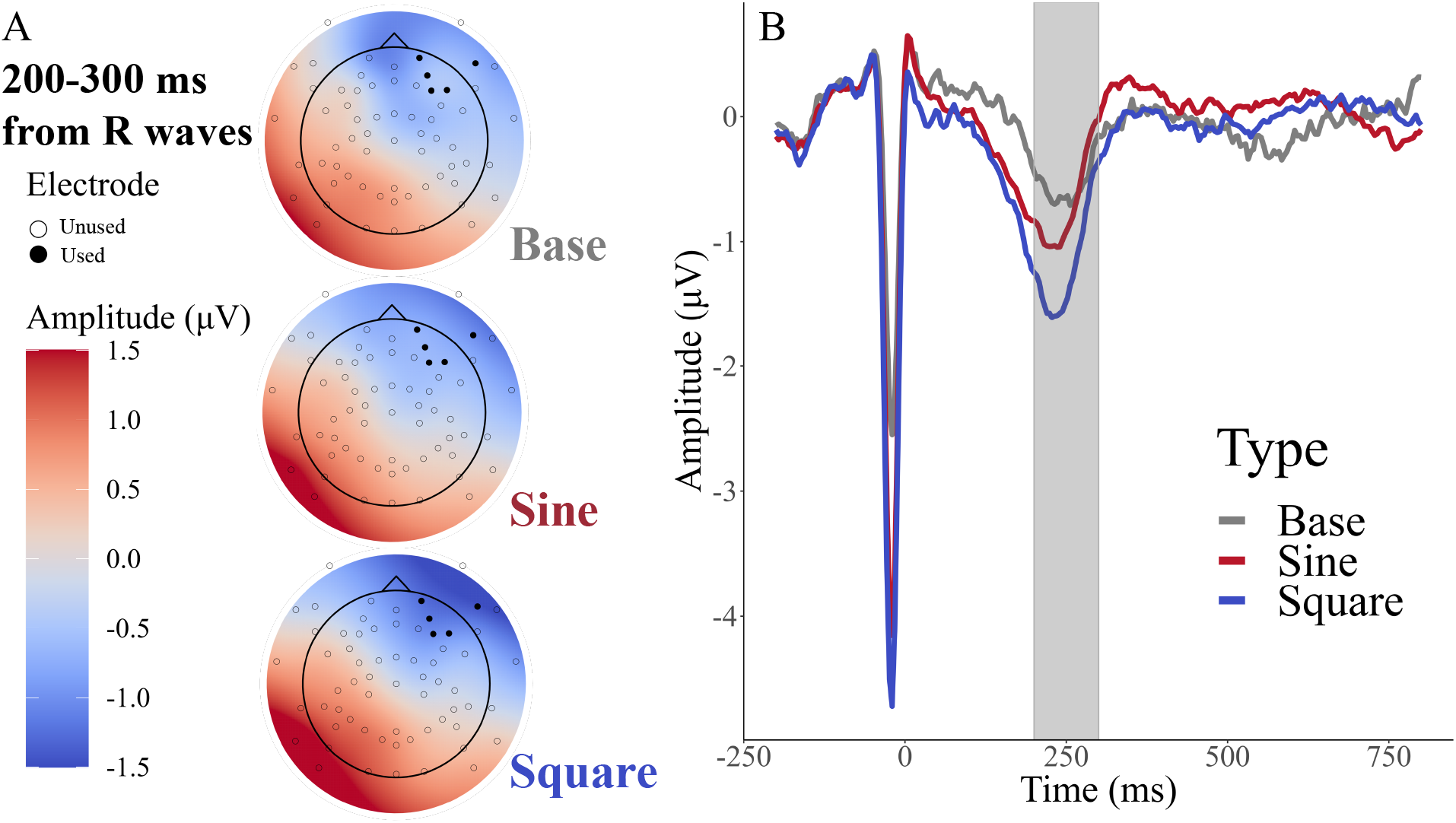
(A) Topographic maps showing the mean voltage of HEP 200–300 ms after the R-wave in each block. The topography maps showed that the amplification of voltage in the negative direction occurred in the right frontal region at approximately 200–300 ms after the R-wave; therefore, the HEP amplitudes of the right frontal electrodes (E1, E2, E5, E59, and E60; indicated by black dots on the topography maps) were averaged for each stimulus type. (B) Average waveforms of the right frontal HEP from R-wave -200–800 ms for each stimulus type (base, sine, and square). The vertical axis shows the amplitude of the EEG and the horizontal axis shows the time from the R-wave of the ECG. The color of the graph indicates the stimulus type, and the light gray bar indicates the time window used to calculate HEP.

### 2.7 Model-Based Analysis

We constructed a generalized mixed model with right frontal HEP amplitude as the objective variable. First, the model included the following five main effects: stimulus type (base, sine, and square), difference in the RR interval, ECG electrode voltage (ECG_amplitude), sex (male or female), and electrical stimulus threshold (threshold). ECG_amplitude, sex, and threshold were included to exclude their effects, as they could have influenced HEP values. In addition, an interaction between stimulus type and the difference in RR intervals and that between the difference in RR intervals and the HCT score were included in the model. The former was used to test the effect of different heart rate changes for each stimulus type on the right frontal HEP amplitude. In the present analysis, we focused on this interaction parameter. The latter was used to determine whether the HEP amplitude is greater in individuals with more accurate interoception, who might be able to perceive the heart rate changes caused by stimulus presentation more easily in their consciousness.

In previous analyses, RR0 was used as the reference point for RR interval in the absence of stimulus presentation and RR1-RR0 and RR2-RR0 were used as measures of changes in heart rate due to stimulus presentation. However, as HEP is the ERP corresponding to one R-wave, we considered it inappropriate to correspond to RR2-RR0, the difference between RR intervals two time points apart. Therefore, in the present model analysis, we used RR2-RR1 instead of RR2-RR0, in addition to RR1-RR0. To examine which of these two RR interval differences better explains the magnitude of the right frontal HEP amplitude, we prepared two models including RR1-RR0 and RR2-RR1 in the RR interval difference part of the model parameters described above, respectively. The two models were compared using the Widely Applicable Information Criterion (WAIC; S. Watanabe, 2010), which estimates the expected log pointwise predictive density for a new dataset that integrates the posterior distributions of the model parameters, allowing for a combined assessment of model fit and complexity. As a lower WAIC value indicates that the model fits the observed data better, a model with a lower WAIC value was adopted.

We used the Bayesian approach to estimate the parameters of the statistical model. This is because we wanted to stabilize the convergence when estimating models with many parameters. The brms R package was used for the analysis (Bürkner, 2017, 2018) and the parameters were estimated using four Markov Chain Monte Carlo chains. Each chain contained 1000 burn-in samples and 2000 additional samples with a thinning parameter of 1, resulting in 1000 posterior samples per chain, which were combined into one posterior sample consisting of 4,000 samples for each parameter. To account for individual differences in the parameters, we employed a model that included the random effects of individual differences in addition to the intercept. The objective variable was assumed to follow a Gaussian distribution. The regression weights had a Cauchy prior distribution centered around zero, with a scale of 1.563. This scale was the same as the SD of the independent variables (Rouder & Morey, 2012). The prior distribution of the SD for group-level effects of the model followed Student’s *t*-distribution around zero, degrees of freedom of 3, and a scale of 2.5. The model intercept, its SD, random effects, intercept of random effects, and residuals of the response variable had uninformed prior distributions. Model convergence was evaluated based on the Gelman–Rubin convergence statistic 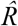 (Gelman & Rubin, 1992), with values close to 1 indicating negligible differences between within- and between-chain variances. The mean and 95% credible interval (CI) were reported as equally-tailed intervals to describe the posterior distributions of sampled regression weights.

## 3. Results

### 3.1 Assessing Group-Level Heart Rate Change Effects: Estimation from the Difference in RR Intervals Before and After Electrical Stimuli

To examine the effect of electrical stimuli on cardiac activity, the mean differences between RR0 and RR1 and RR2 (i.e., RR1-RR0 and RR2-RR0) were calculated for each condition (Fig. 5). Significant differences were observed between RR1-RR0 and 0 at the 1% level for sine_50 and square_50 and at the 5% level for sine_90 (sine_50: *t* (41) = 3.983, *p* = .001, *d* = .463; sine_90: *t* (42) = 2.388, *p* = .036, *d* = .305; square_50: *t* (41) = 3.479, *p* = .003, *d* = .463). In contrast, there was no significant difference between the average RR interval and 0 for base and square_90 of RR1-RR0 and all stimulus conditions of RR2-RR0 (RR1-RR0; base: *t* (42) = -1.030, *p* = .386, *d* = .000; square_90: *t* (42) = 1.275, *p* = .262, *d* = .152; RR2-RR0; base: *t* (42) = .504, *p* = .617, *d* = .152; sine_50: *t* (42) = 2.039, *p* = .208, *d* = .305; sine_90: *t* (42) =2.371, *p* = .112, *d* = .305; square_50: *t* (42) = 1.566, *p* = .208, *d* = .152; square_90: *t* (42) = 0.395, *p* = .395, *d* = .000). For sine_50, sine_90, and square_50, the RR interval tended to increase at RR1; that is, the heart rate tended to decrease immediately after the presentation of the subthreshold electrical stimuli.

**Fig. 5.**
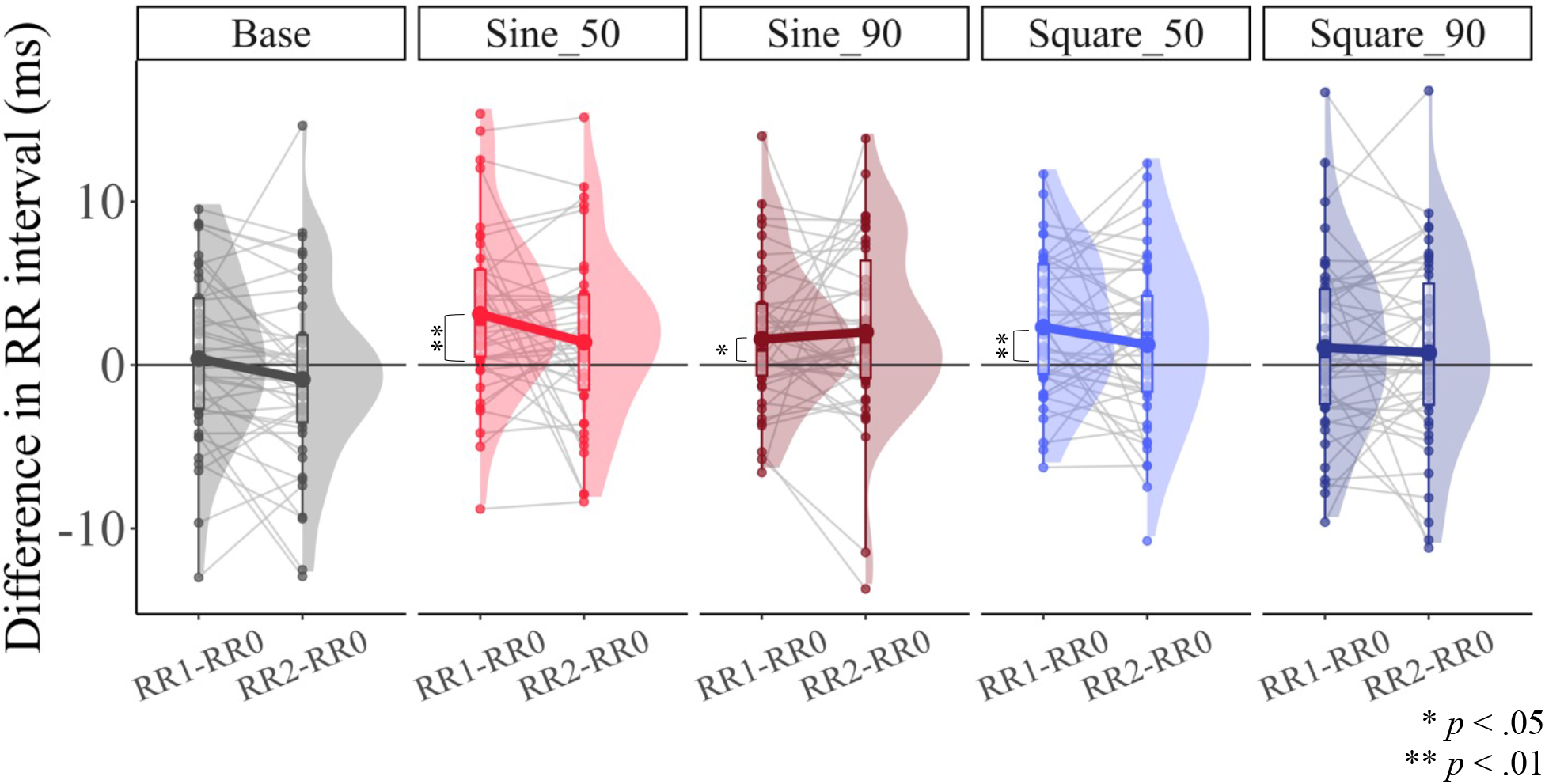
Distribution of RR1-RR0 (difference between the RR interval of the first beat after stimulus presentation and that before stimulus presentation) and RR2-RR0 (difference between the RR interval of the second beat after stimulus presentation and that before stimulus presentation) for each condition. In the base condition, no subthreshold current stimuli were presented. Sine wave stimuli were presented in the sine_50 and sine_90 conditions, and square wave stimuli were presented in the square_50 and square_90 conditions with an intensity of 50% and 90% of the participant’s threshold, respectively. The density plots show the distribution of RR intervals per condition; the box plot shows the minimum and maximum values, first, second, and third quartiles of RR interval data per condition; and the points below the box plot show the actual RR interval per condition for the participants. If the difference is greater than 0, the value of the RR interval at each time point decelerated from RR0; if it is less than 0, it accelerated from RR0. If the line on the graph is down to the right, the heart rate accelerated from RR1 to RR2. If the line is up to the right, the heart rate decelerates from RR1 to RR2. Only RR1-RR0 for sine_50, sine_90, and square_50 showed a statistically significant difference between 0. The distribution of heart rate data indicated that these conditions caused a decrease in heart rate immediately after the presentation of the subthreshold stimuli.

In addition, the repeated measures ANOVA on the means of each block of RR1-RR0 and RR2-RR0 showed no interaction or main effect of stimulus intensity and type for either RR1-RR0 or RR2-RR0 (RR1-RR0; type: *F* (1, 166) = .765, *p* = .383, *η*^2^ = .005; intensity: *F* (1, 166) = 3.541, *p* = .062, *η*^2^ = .021; type*intensity (interaction between type and intensity, hereafter interactions are expressed using *): *F* (1, 166) = .780, *p* = .855, *η*^2^ = .000; RR2-RR0; type: *F* (1, 167) = .720, *p* = .397, *η*^2^ = .004; intensity: *F* (1, 167) = .006, *p* = 0.938, *η*^2^ = .000; type*intensity: *F* (1, 167) = .431, *p* = .512, *η*^2^ = .003).

### 3.2 Relationship Between Changes in Peripheral Cardiac Activity Induced by Electrical Stimuli and Afferent Signal Processing

To investigate whether the difference in RR intervals immediately after stimulus presentation and the stimulus type affected the amplitude of the right frontal HEP, we performed the modeling described in section 2.6. After comparing the models using WAIC, we adopted a model in which RR2-RR1 was input as a parameter of the difference in the RR interval (WAIC: RR1-RR0, 422.0; RR2-RR1, 412.4; the estimated values of each parameter in the model adopting RR1-RR0 are shown in Appendix Table A.1). Table 1 and Fig.6 summarize the estimated coefficients for the parameters that are important for the interpretation and discussion of this model. As indicated by 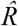 equal to 1, all chains converged to the model.

**Table 1.**
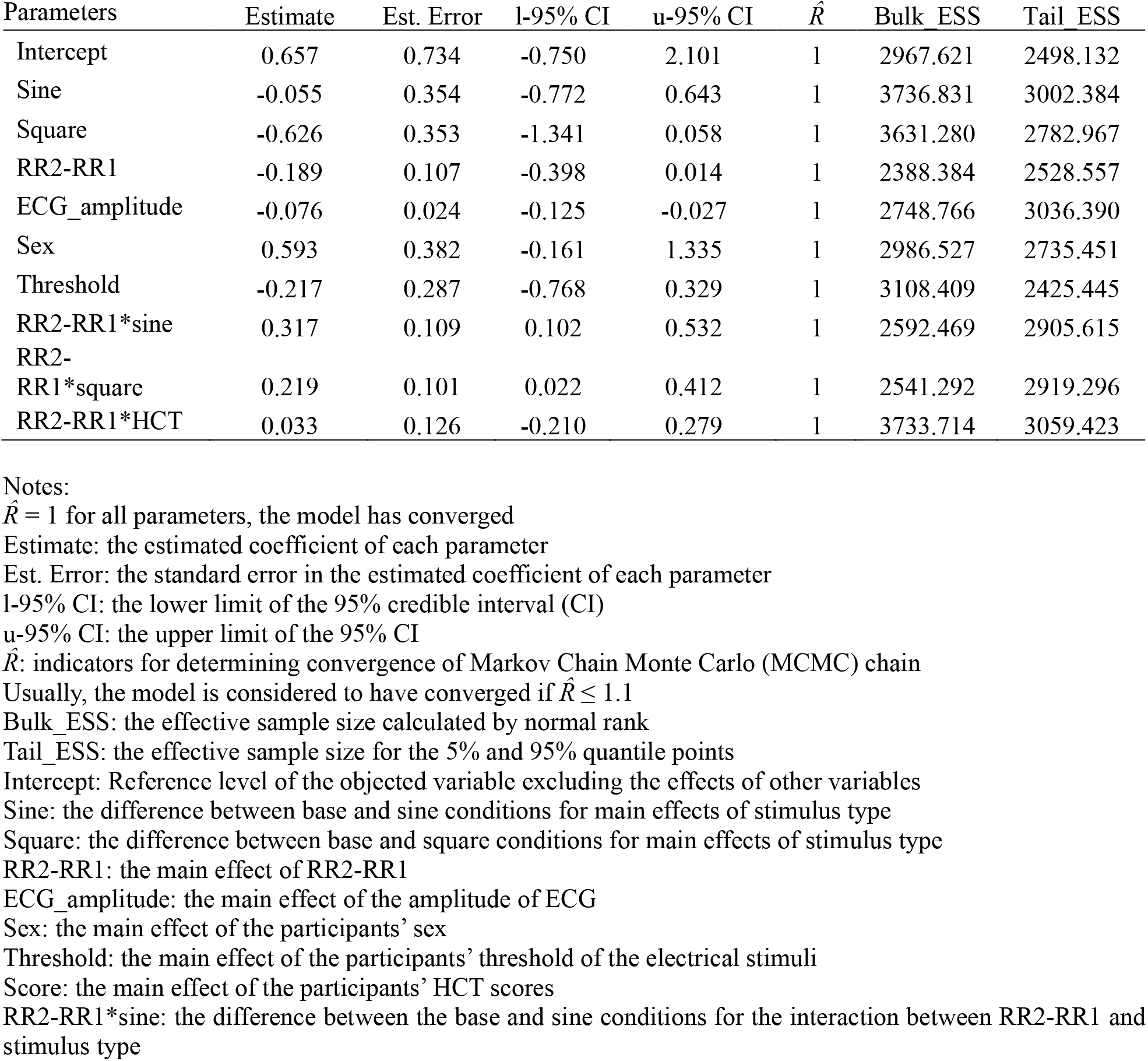
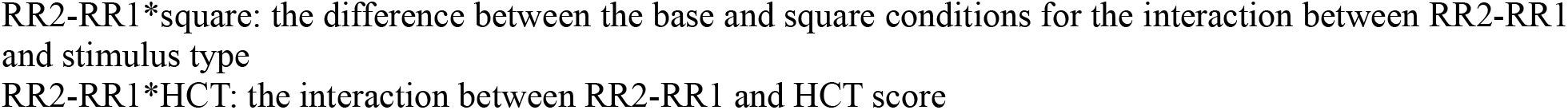
Summary of the estimated parameters.

| Parameters | Estimate | Est. Error | l-95% CI | u-95% CI | $\hat{R}$ | Bulk ESS | Tail ESS |
| --- | --- | --- | --- | --- | --- | --- | --- |
| Intercept | 0.657 | 0.734 | -0.750 | 2.101 | 1 | 2967.621 | 2498.132 |
| Sine | -0.055 | 0.354 | -0.772 | 0.643 | 1 | 3736.831 | 3002.384 |
| Square | -0.626 | 0.353 | -1.341 | 0.058 | 1 | 3631.280 | 2782.967 |
| RR2-RR1 | -0.189 | 0.107 | -0.398 | 0.014 | 1 | 2388.384 | 2528.557 |
| ECG_amplitude | -0.076 | 0.024 | -0.125 | -0.027 | 1 | 2748.766 | 3036.390 |
| Sex | 0.593 | 0.382 | -0.161 | 1.335 | 1 | 2986.527 | 2735.451 |
| Threshold | -0.217 | 0.287 | -0.768 | 0.329 | 1 | 3108.409 | 2425.445 |
| RR2-RR1*sine | 0.317 | 0.109 | 0.102 | 0.532 | 1 | 2592.469 | 2905.615 |
| RR2-RR1*square | 0.219 | 0.101 | 0.022 | 0.412 | 1 | 2541.292 | 2919.296 |
| RR2-RR1*HCT | 0.033 | 0.126 | -0.210 | 0.279 | 1 | 3733.714 | 3059.423 |

Notes:
$\hat{R} = 1$ for all parameters, the model has converged
Estimate: the estimated coefficient of each parameter
Est. Error: the standard error in the estimated coefficient of each parameter
l-95% CI: the lower limit of the 95% credible interval (CI)
u-95% CI: the upper limit of the 95% CI
$\hat{R}$ : indicators for determining convergence of Markov Chain Monte Carlo (MCMC) chain
Usually, the model is considered to have converged if $\hat{R} \leq 1.1$
Bulk ESS: the effective sample size calculated by normal rank
Tail ESS: the effective sample size for the 5% and 95% quantile points

The posterior predicted values of HEP for each stimulus type are shown in Fig. 7A. Although there was no main effect of stimulus type (Table 1), the absolute value of the coefficient estimate was larger in the square-wave condition than in the sine-wave condition (*b*_*sine*_ = -.055; *b*_*square*_ = -.626). Compared with the base condition, the posterior predictive value of the right frontal HEP in the square-wave condition was negatively amplified. In addition, there was an interaction between RR2-RR1 and type for both sine and square conditions (Fig. 6; sine: *b* = .317 95%CI [.102;.532]; square: *b* = .219 95%CI [.022;.412]). The relationship between RR interval and HEP was reversed under the base and stimulus presentation conditions (Fig. 7B). The smaller the value of RR2-RR1 in both the sine and square conditions, that is, the greater the heartbeat acceleration in RR2 compared with RR1, the greater the amplitude of the right frontal HEP. This effect was larger in the sine condition than in the square condition (*b*_*RR2-RR1_sine*_ = .317; *b*_*RR2-RR1_square*_ = .219). Although there was a main effect of the ECG_amplitude (Table 1, Fig. 6), which is an artifact relative to the HEP signal, the estimated value of the coefficient was smaller than that of the other effects (*b* = -.076, 95%CI [-.125;-.027]).

**Fig. 6.**
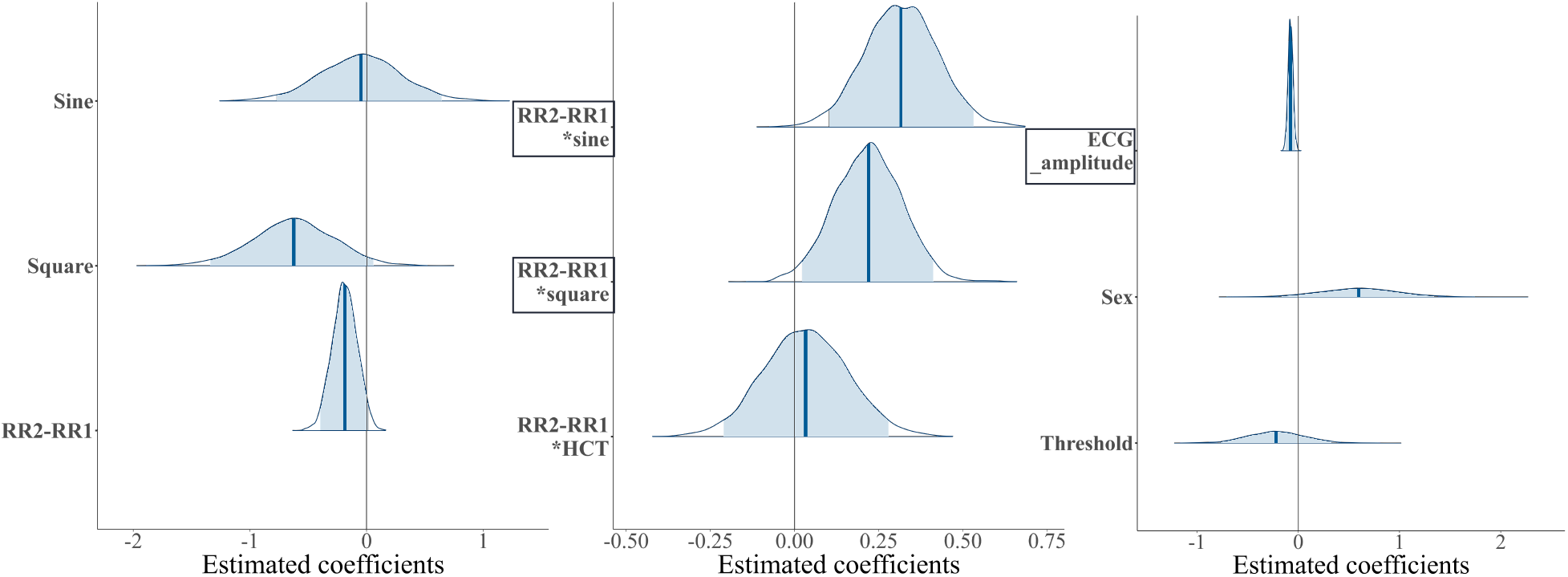
Distribution of the estimated parameters. The horizontal axis represents the estimated coefficients for each factor. Dark blue lines indicate the mean of the estimated coefficients and the light blue ranges indicate 95% credible intervals (CI). Parameter names in bold boxes indicate that the 95% CI was not across 0. The interaction between the difference in RR intervals and each stimulus type (sine/square) and the main effect of the ECG voltage magnitude influenced the value of the right frontal HEP amplitude.

**Fig. 7.**
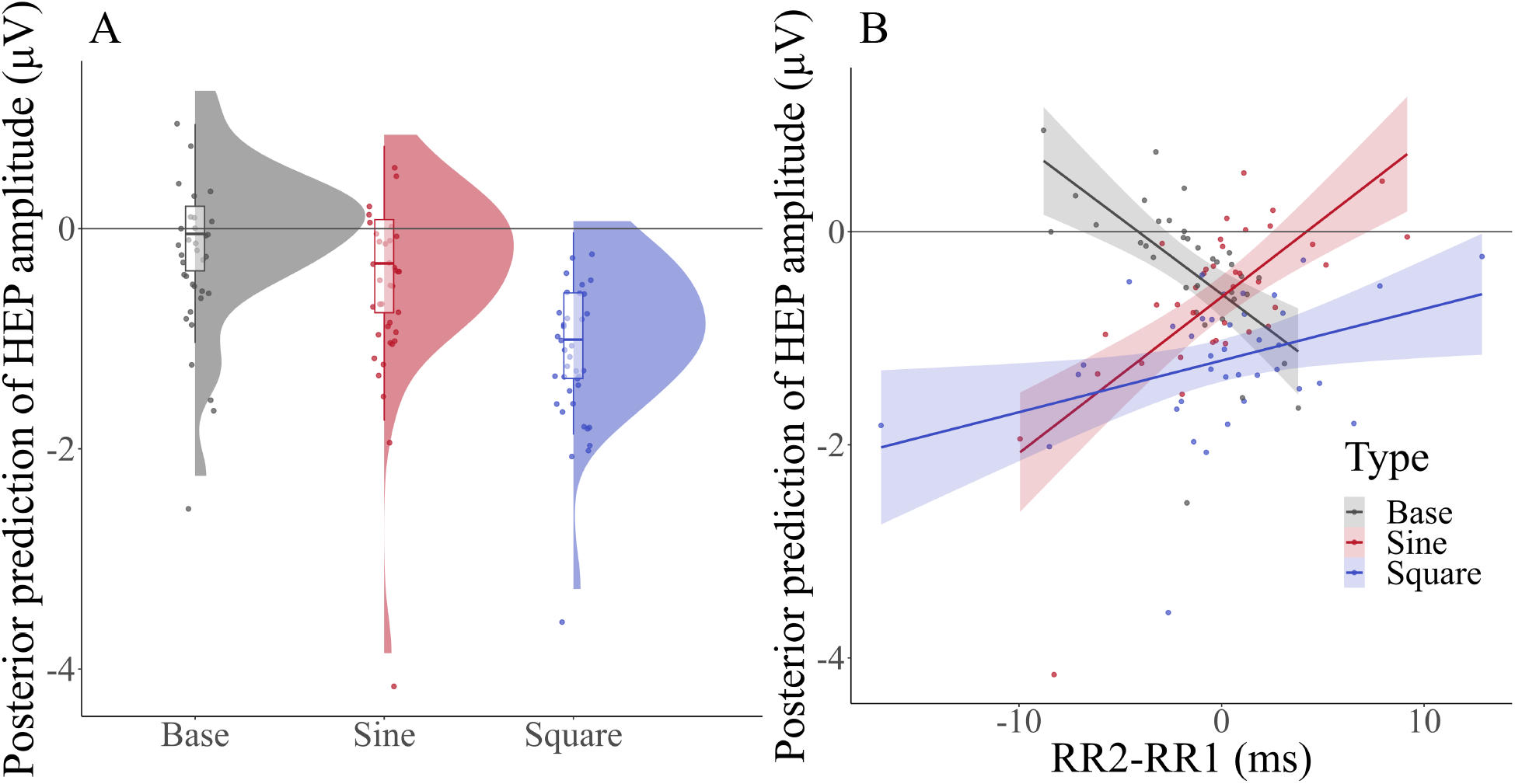
(A) Posterior prediction of right frontal HEP amplitude for each stimulus type (base/sine/square). The density plots show the distribution of the predictive amplitude; the box plot shows the minimum and maximum values, the first, second, and third quartiles of the predicted amplitude; and the points below the box plot show the predicted amplitudes obtained at each sampling. Although there was no main effect of stimulus type, the posterior predicted amplitude of the right frontal HEP in the square condition tended to amplify in a negative direction than in the other two conditions. (B) Posterior predictive distribution of the relationship between the stimulus type and RR2-RR1. The horizontal axis shows the RR2-RR1 and the vertical axis shows the estimated value of the HEP amplitude. The solid line in the graph represents the regression line based on estimated values.

Each colored range indicates the 95% CI and the dots indicate the posterior predicted values for each participant. In the sine/square condition, the smaller the value of RR2-RR1, that is, the greater the acceleration of the heartbeat, the more the amplification of HEP in the negative direction, and this effect was greater in the sine condition.

## 4. Discussion

This study investigated how subthreshold tactile stimuli affect ANS activity and afferent signal processing. We used heart rate as a measure of ANS activity and HEP for afferent signal processing.

The analysis showed that heart rate decelerated at RR1 for sine_50, sine_90, and square_50 (Fig. 5). However, this effect was not observed at RR1 in the base and square_90 and at RR2 in all conditions. The Bayesian generalized mixed model with the right frontal HEP amplitude as the objective variable showed that the right frontal HEP amplified in a negative direction as the heartbeat accelerated in RR2 in the condition in which subthreshold electrical stimuli were presented (sine/square). This effect was stronger under the sine condition (Fig. 7B).

The results partially supported hypothesis (1), as heart rate decelerated immediately after the presentation of subthreshold tactile stimuli in the three conditions of sine_50, sine_90, and square_50. As no such tendency was observed in the base condition without stimulus presentation, the deceleration of the heartbeat observed in the stimulus conditions was considered more pronounced than the fluctuations normally associated with human heartbeats. However, there was no heart rate change effect in square_90, despite the higher stimulus intensity. Moreover, in the sine condition, where heart rate change effects were observed at both intensities and the effect size was smaller in the subthreshold 90% condition (see section 3.1). Therefore, hypothesis (2) was not supported. Hypothesis (3) was supported because the HEP amplitude increased as the heart rate accelerated at RR2. The results suggest that even tactile stimuli that are not consciously perceived can influence ANS activity and afferent signal processing.

### 4.1 Changes in Heart Rate in the Orienting Response to Subthreshold Stimuli

Heart rate deceleration after the presentation of the subthreshold stimuli could be due to changes in ANS activity associated with the orienting response to the stimulus (Graham & Clifton, 1966; Simons et al., 1998). This phenomenon has been confirmed by the presentation of suprathreshold visual, auditory, and tactile stimuli (Graham & Clifton, 1966; Grund et al., 2022; Motyka et al., 2019; Stekelenburg & van Boxtel, 2002). It is an automatic, pre-attentive change in brain activity and peripheral reflex response to unexpected and novel changes in the environment (Sokolov, 1963). The occurrence of the orienting response and accompanying changes in ANS activity may not require suprathreshold perception of the stimulus. Orienting responses and associated changes in the ANS to subthreshold visual stimuli have been reported (Iida et al., 2012; Maoz et al., 2012; Ruiz-Padial et al., 2005; Sebastiani et al., 2011; van der Ploeg et al., 2017, 2020). To the best of our knowledge, no studies have directly examined orienting responses to subthreshold tactile stimuli. Information about the subthreshold tactile stimuli reaches the primary somatosensory cortex and the ERP components, P50 or P60, are observed after their presentation (Forschack et al., 2017; Iliopoulos et al., 2020; Nierhaus et al., 2015). This suggests that even tactile stimuli that cannot be consciously perceived may elicit orienting responses during such information processing.

### 4.2 Differences in Heart Rate Change Effects by Stimulus Type and Intensity

The present study found no significant main effect of stimulus intensity on the difference in RR intervals (see Section 3.1), nor any significant correlation between participants’ stimulus thresholds, that is, the physical intensity of the stimuli presented during the task and the difference in RR intervals (Appendix Fig. A.3). These results suggest that the physical intensity of the subthreshold tactile stimulus does not affect the degree of the heart rate change. Based on the results of our previous studies using subthreshold tactile stimuli, we hypothesized that a higher stimulus intensity would have a greater effect on heart rate, even in subthreshold tactile stimuli; however, this hypothesis was not supported. The results of this study do not reveal whether subthreshold tactile stimuli have a specific relationship with ANS activity, different from that of suprathreshold stimuli, or the differences in intensity between subthreshold stimuli are so small that they do not appear as differences in heart rate changes. However, an individual’s subjective tactile threshold is likely to be different from the stimulus intensity at which subthreshold tactile stimuli affect the ANS activity. The two intensities should not be considered identical in future studies.

It is noteworthy that the participants’ heart rate significantly changed only in the sine wave condition, regardless of stimulus intensity. In the sine wave condition, the effect size of the heart rate changes was greater in the 50% condition than in the 90% condition (*d*_*sine_50*_ = .463 > *d*_*sine_90*_ = .305). By contrast, in the square wave condition, the heart rate change effect was observed only in the 50% threshold condition (Fig. 5).

Although the interaction between stimulus type and stimulus intensity was not statistically significant (see section 3.1), these results suggest that even subthreshold tactile stimuli presented to the same body part may have different optimal intensities to induce heart rate changes, depending on the stimulus type. For suprathreshold stimuli, the heart rate deceleration, which is part of the orienting response, intensifies as stimulus intensity is reduced and detectability of the stimuli decreases (Graham, 1979; Jackson, 1974; Turpin et al., 1999). Differences in saliency between stimulus types may also affect the detectability of suprathreshold stimuli. Unlike square waves, which are instantaneous, discrete, and sharp, sine waves are continuous and gradually spread as waves. A previous study showed that square waves are easier to detect than sine waves when a suprathreshold electrical stimulus is applied to the fingertip at frequencies below 60 Hz (Vardar et al., 2017). If a similar phenomenon occurred for subthreshold tactile stimuli, ambiguous sine waves would be more likely than square waves to produce a significant decrease in the heart rate, regardless of the stimulus intensity. Conversely, a more distinct stimulus, the square wave, would be less likely to cause a heart rate deceleration, causing no significant change in the 90% threshold condition. In the future, the relationship between the differences in the saliency of tactile stimuli and changes in ANS activity needs to be examined using suprathreshold tactile stimuli, considering the characteristics of the receptors on the skin and other factors.

### 4.3 Afferent Signal Processing of Heart Rate Changes and Implications for HEP Research

In this study, right frontal HEP amplitudes tended to amplify more in the sine and square wave conditions in which subthreshold electrical stimuli were presented, than in the base condition without them (Fig. 7A). In addition, the amplitude of the right frontal HEP was greater in participants whose heart rate accelerated to a greater degree in the second beat after the stimulus presentation (Fig. 7B). As described in Section 3.2, the model including RR1-RR0 had less explanatory power than that including RR2-RR1. Moreover, the coefficients of the interaction between RR interval difference and stimulus type in the model including RR1-RR0 were *b*_*RR1-RR0_sine*_ = -.013 95%CI [-.219;.193] and *b*_*RR1-RR0_square*_ = .036 95%CI [-.134;.207], indicating that this interaction was ineffective (See Appendix Table A.1). These results indicate that the acceleration of the heart rate on the second beat after stimulus presentation specifically influenced the difference in HEP amplitude across blocks, rather than the deceleration on the first beat.

The effects of heart rate changes on HEP differed between the time points from stimulus presentation, which may be due to stronger autonomic regulation on the second beat after stimulus presentation. In the orienting response to suprathreshold stimuli, heart rate deceleration followed by acceleration has been observed (Graham, 1979 Smith & Strawbridge, 1969; Wood & Obrist, 1964). This acceleration may be a defensive response to nociceptive stimuli (Graham, 1979; Velden & Schumacher, 1979) or loss of vagal tone during the inspiratory phase of breathing (Smith & Strawbridge, 1969; Wood & Obrist, 1964). As the tactile stimuli used in this study were below the threshold, it is unlikely that the participants felt threatened by the stimuli and that a defensive response was generated. It is not yet clear whether the latter factor is an incidental change due to the respiratory cycle unrelated to stimulus presentation or the result of autonomous regulation to maintain homeostasis in response to a decrease in the heart rate. However, the second beat after subthreshold tactile stimulation, the heart rate is accelerated by respiration and this increased information about the activity of ANS may amplify the HEP. If such modulation of respiration is an autonomic adjustment for homeostasis, it may explain why the interaction effect between post-stimulus heart rate acceleration and HEP amplitude increase was greater in the sine wave condition. As noted in Section 4.2, sine wave stimuli were significantly more salient than square waves and more strongly decelerated heart rate. This greater deceleration may also more strongly activate the ability to restore heart rate levels by breathing. In the future, respiration should also be measured to gain an integrated understanding of the causal relationship between subthreshold tactile stimuli, changes in heart rate, and increases in HEP amplitude.

The results of the present study suggest that a bottom-up ECG input with changes in peripheral heart rate increases the amplitude of HEP. This is important in interpreting the information reflected by HEP. Many previous studies have focused on HEP measured during cognitive activities involving direct attention to the body such as heartbeat detection (Katkin et al., 1991; Montoya et al., 1993; Petzschner et al., 2019; Pollatos & Schandry, 2004; Schandry & Weitkunat, 1990; Yuan et al., 2008). The relationships among cognitive activities, such as emotion (Couto et al., 2015; MacKinnon et al., 2013), pain (Shao et al., 2011), arousal (Luft & Bhattacharya, 2015), time perception (Richter & Ibáñez, 2021), and self-referential processing (Babo-Rebelo, Richter, et al., 2016; Babo-Rebelo, Wolpert, et al., 2016; Huang et al., 2023), which are potentially influenced by body signal processing, have also been the focus of attention. Few studies have addressed the direct relationship between peripheral cardiac activity parameters and HEP modulation, suggesting that mechanoreceptors in the cardiac wall, baroreceptors in the aortic arch, and subsequent RR interval changes are associated with HEP (Baranauskas et al., 2021; Gray et al., 2007; Park et al., 2018). In the present study, participants were not instructed to pay attention to bodily sensations; therefore, they knew neither the stimuli to elicit heart rate change nor the change itself. However, the more accelerated the heartbeat in RR2, the more amplified was the right frontal HEP amplitude (Fig. 7B). This effect was sufficiently large (Table 1), even after excluding the effect of the ECG electrode voltage, which has been treated as an artifact in previous HEP studies. The fact that the interaction between RR2-RR1 and HCT performance did not affect the HEP amplitude may be important for understanding interoception. This result suggests that high or low interoceptive accuracy, as measured by HCT, may not correlate with the degree of afferent signal transmission associated with peripheral heart rate changes. Perhaps interoceptive accuracy may indicate the extent to which HEP influences cognitive activity related to somatosensation, as described above. Thus, methods that can induce changes in peripheral heart rate without attracting the participant’s attention, such as those used in this study, have the potential to refine models of interoception and related cognitive activity.

### 4.4 Limitation

This study had two limitations. First, subthreshold electrical stimuli caused significant noise in the EEG; therefore, epochs corresponding to RR1 were often excluded from the analysis. This may explain why the analysis of the model with right frontal HEP amplitude, including the RR2-RR1 values, was evaluated as more valid than that of the model that included the RR1-RR0 values of the ECG. We are currently investigating the use of a device that minimizes noise in the physiological index data being measured in parallel. In the future, this device may allow the calculation of relevant ERP and HEP amplitudes at each time point after stimulus presentation and may reveal the time-series causal relationship between the processing of subthreshold tactile stimuli during unconsciousness and subsequent heart rate changes and their signal processing. Second, the extent to which the heart rate change effect of the subthreshold tactile stimuli observed in this study can be generalized is unclear. In this study, the magnitude of the right frontal HEP amplitude was unaffected by sex (Table 1). However, responsiveness to current stimulation and modulation of HEP may be affected by age (Kamp et al., 2021), body shape, and personality factors, such as anxiety tendency (Domschke et al., 2010; Judah et al., 2018; Verdonk et al., 2024), and the effects of these factors should be examined in the future.

### 4.5 Conclusion

The present study found that the presentation of subthreshold tactile stimuli reduced heart rate immediately after stimulation. The right frontal HEP amplitude was greatly amplified, with greater heart rate changes induced by subthreshold tactile stimulation. These results indicate that changes in ANS activity associated with orienting response can occur even in response to tactile stimuli that are not perceived consciously. Moreover, peripheral heart rate changes can affect the HEP, afferent signal processing of cardiac signals. In the future, it will be necessary to investigate the robustness of the heart rate change effect, observed in this study, with subthreshold tactile stimulation and its effects on somatosensory-related cognitive activities such as emotion, memory, and decision-making.

## Ethics approval

This study was approved by the Keio University Research Ethics Committee (No. 220190100), Japan. The study was conducted in accordance with the 1964 Declaration of Helsinki and its later amendments. All participants provided written informed consent to participate in this study.

## Data Availability Statement

The datasets analyzed in this study are available from the corresponding author upon reasonable request.

## CRediT authorship contribution statement

**Mai Sakuragi:** Conceptualization, Methodology, Software, Formal analysis, Investigation, Writing - Original Draft, Writing - Review & Editing; **Yuto Tanaka:** Methodology, Formal Analysis, Writing - Review & Editing; **Kazushi Shinagawa:** Formal Analysis, Writing - Review & Editing; **Koki Tsuji:** Methodology, Writing - Review & Editing; **Satoshi Umeda:** Conceptualization, Methodology, Formal Analysis, Writing - Review & Editing, Supervision, Funding acquisition.

## Acknowledgments

We are deeply grateful to all the participants of this study. Their willingness to contribute to their time and effort was essential to the success of this study.

