## Supplementary material for "Effects of unconscious tactile stimuli on autonomic nervous activity and afferent signal processing": Fig. A.1.-A.9., Table A.1.

### Appendix

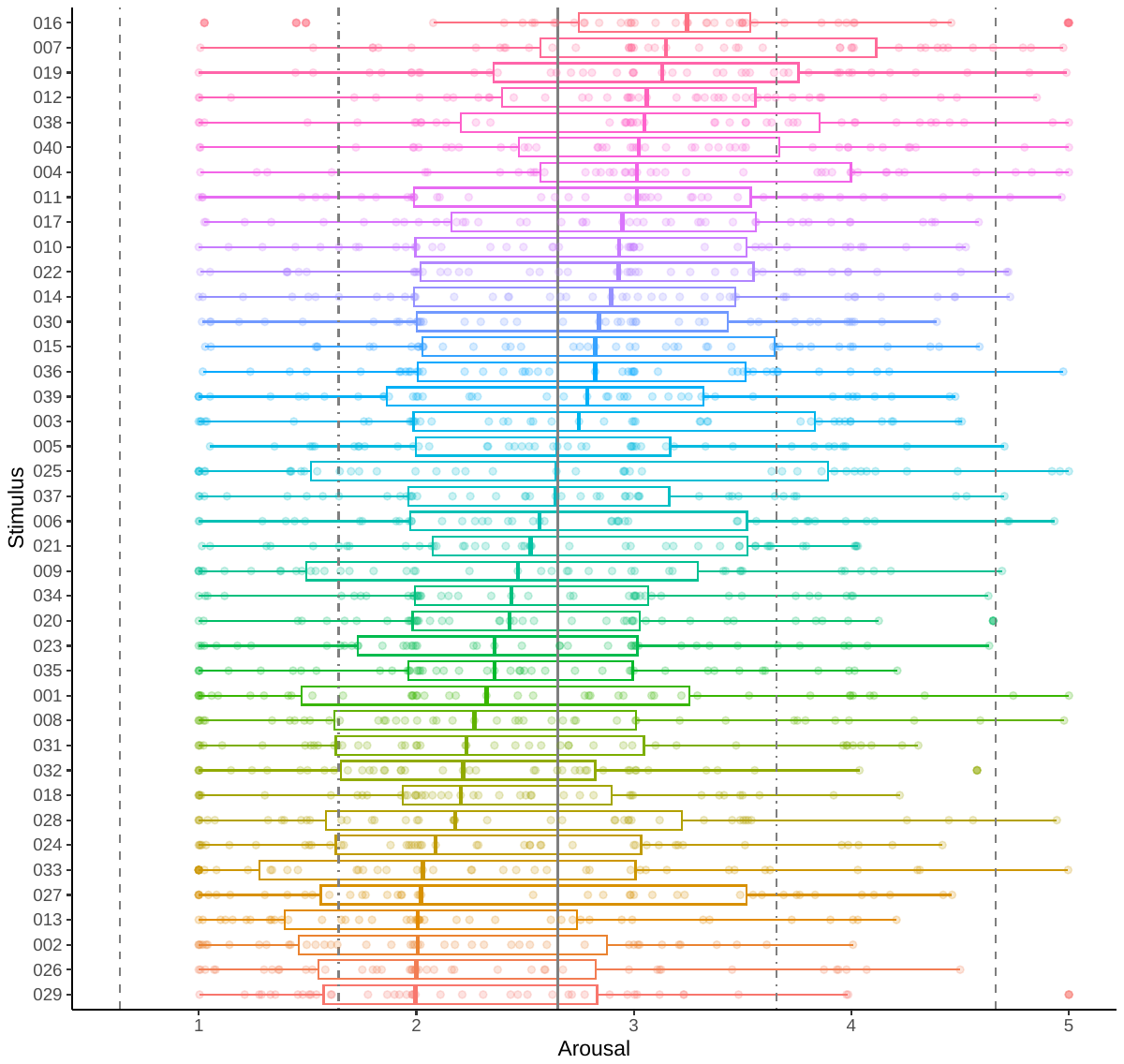

**Fig. A.1.** Distribution of arousal ratings for each stimulus used in the image-viewing task. The box plot shows the minimum and maximum values and the first, second, and third quartiles of arousal ratings, and the points on the box plot show the actual arousal ratings for individuals. The solid line in the center of the graph shows the mean of the evaluated values of arousal for all images, and the two dashed lines on each side show the mean ±1 standard deviation and mean ±2 standard deviation.

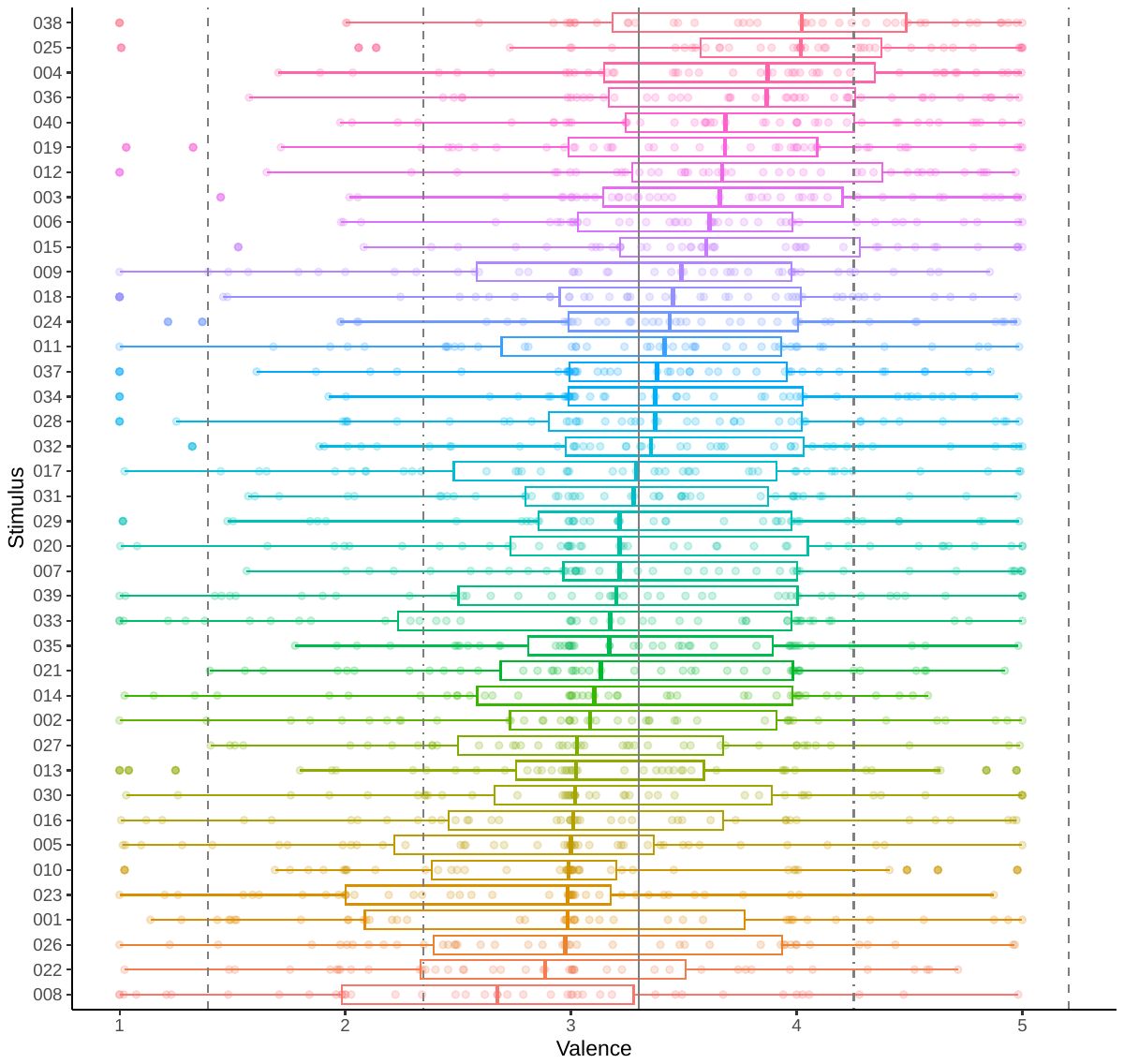

**Fig. A.2.** Distribution of valence ratings for each stimulus used in the image-viewing task. The box plot shows the minimum and maximum values and the first, second, and third quartiles of valence ratings, and the points on the box plot show the actual valence ratings for individuals. The solid line in the center of the graph shows the mean of the evaluated values of valence for all images, and the two dashed lines on each side show the mean ±1 standard deviation and mean ±2 standard deviation.

Fig. A.3. to Fig. A.8. show the correlations between the values of RR1-RR0 and RR2-RR0 for each stimulus type and the participant’s threshold of the electrical stimuli, the scores of the HCT, the HCT's confidence, MAIA's subordinate conceptions, such as noticing, attention regulation, body listening, and BPQ. Colors indicate the stimulus types. The solid line in the graph represents a regression line. Each colored range indicates the 95% CI. The points show the actual value.

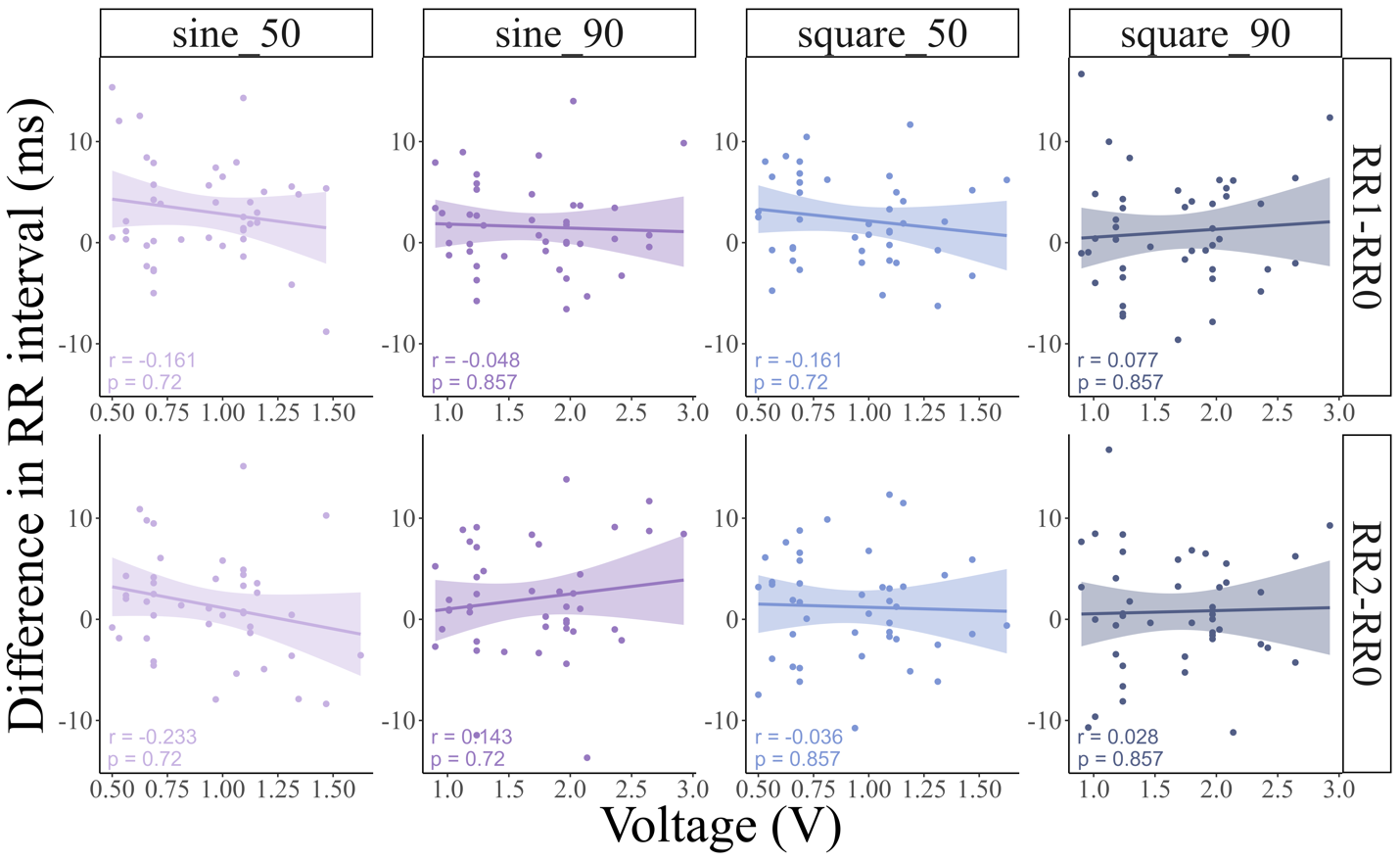

**Fig. A.3.**

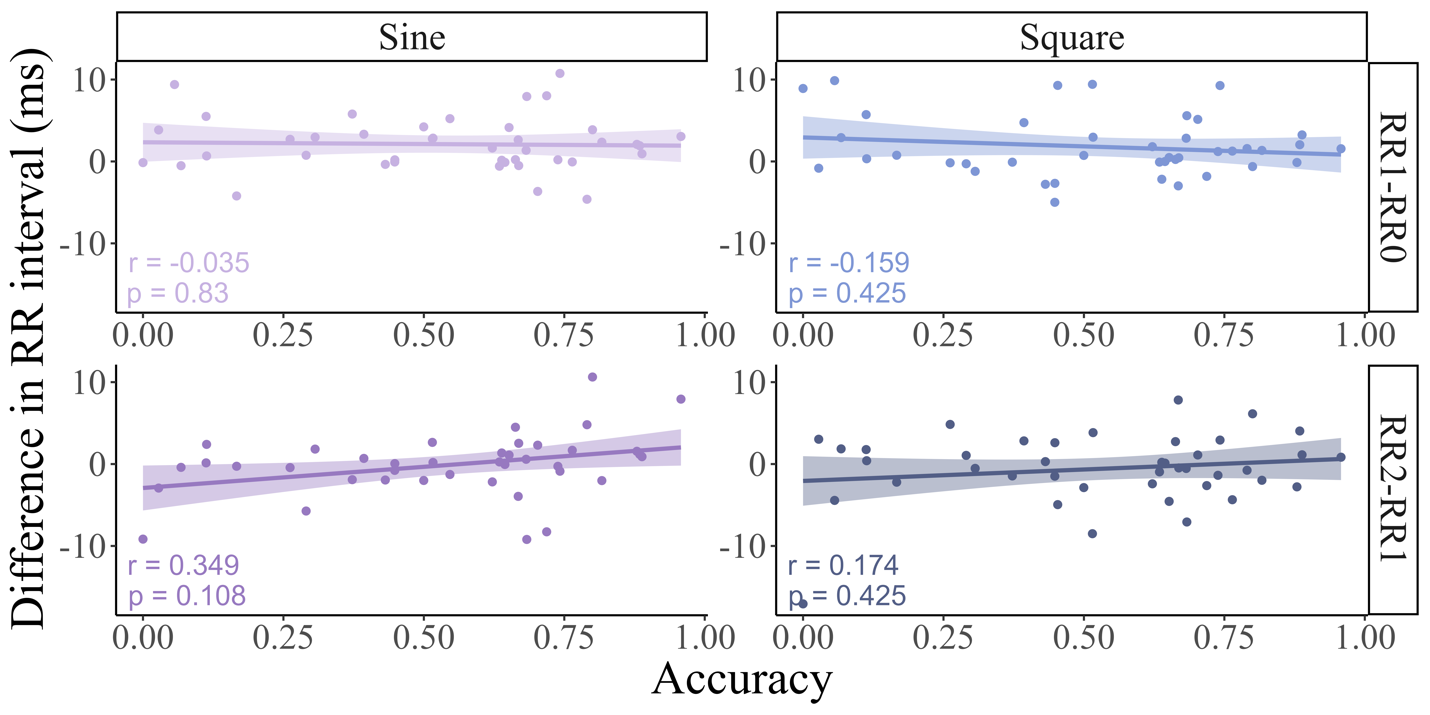

**Fig. A.4.**

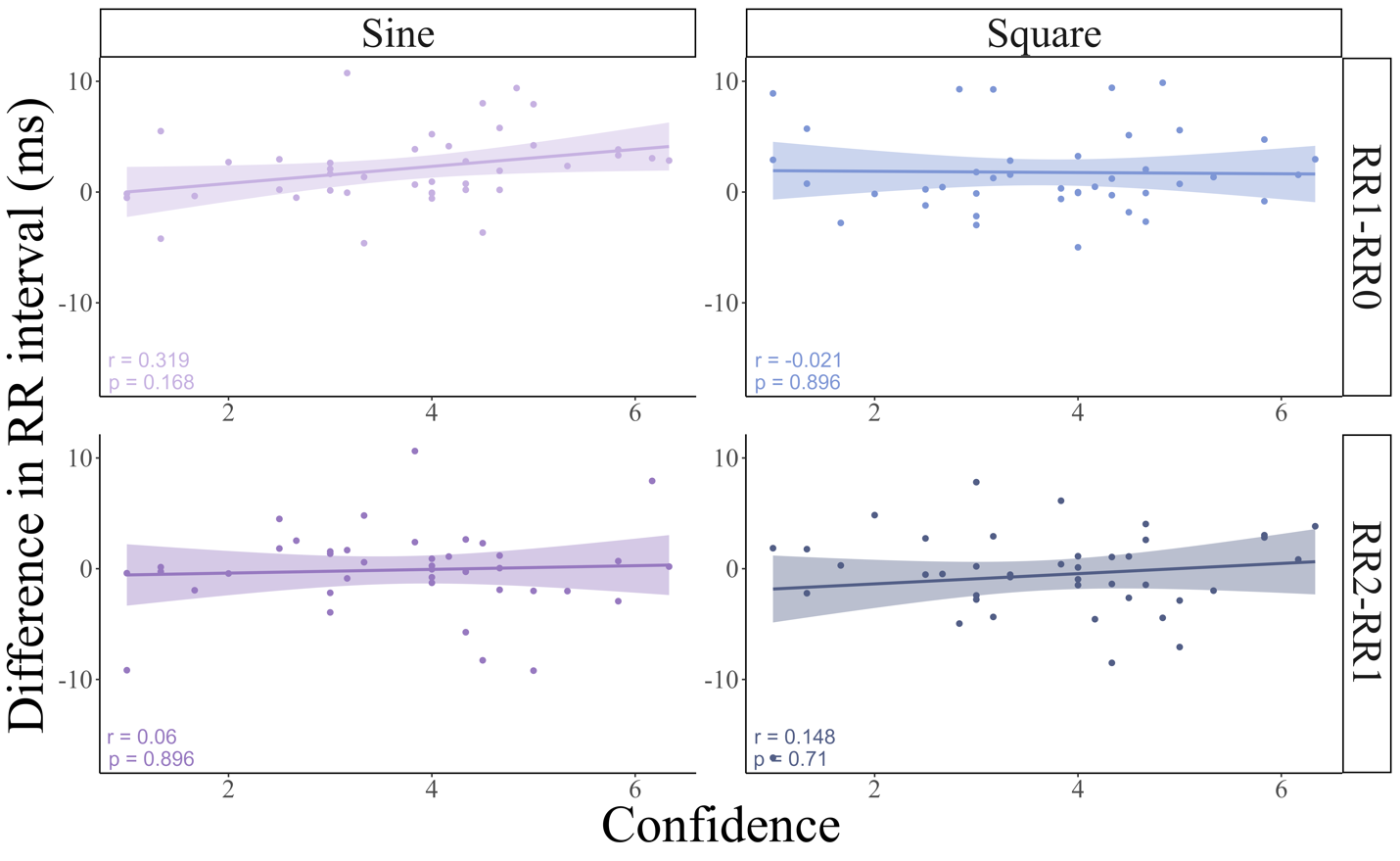

**Fig. A.5.**

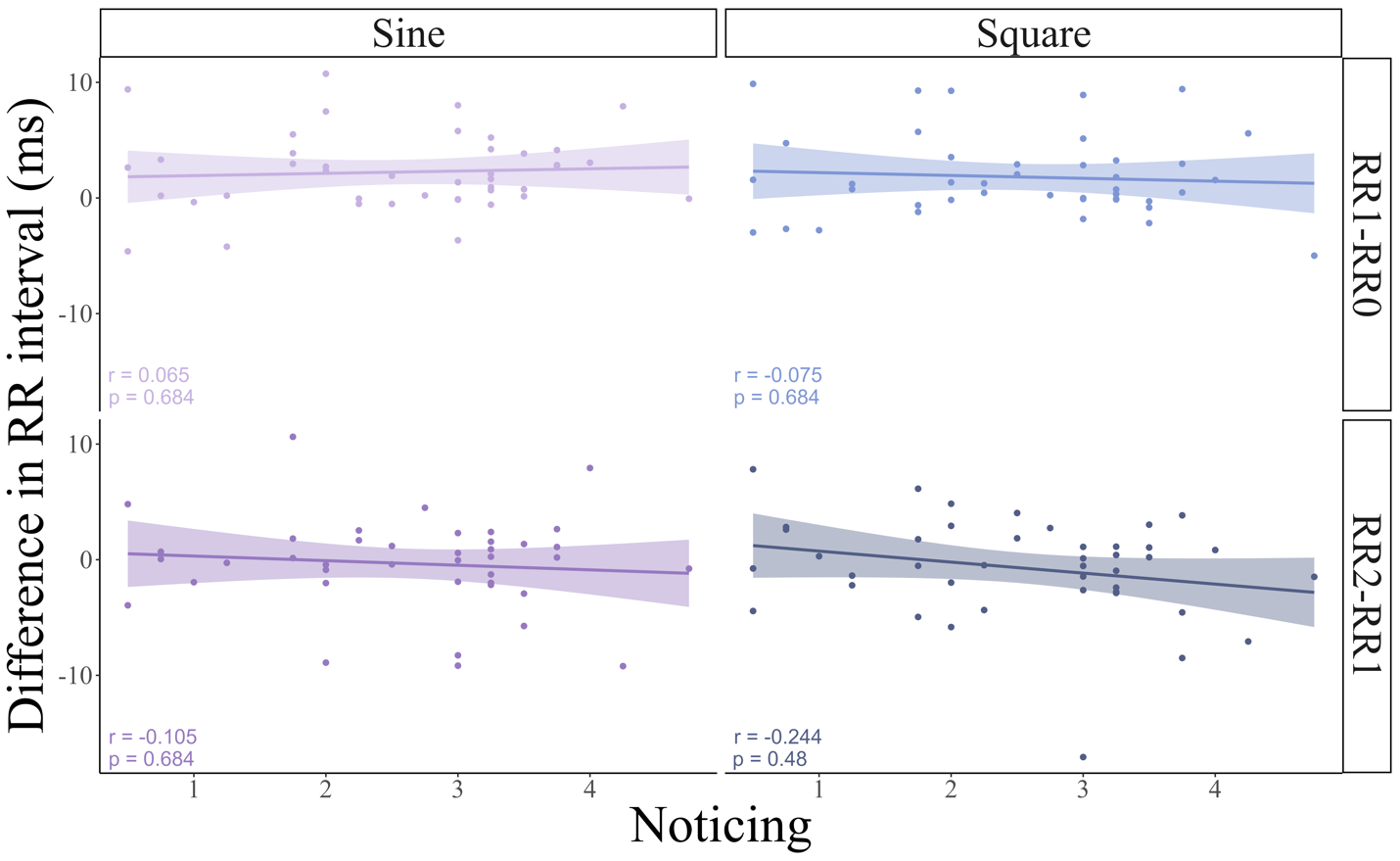

**Fig. A.6.**

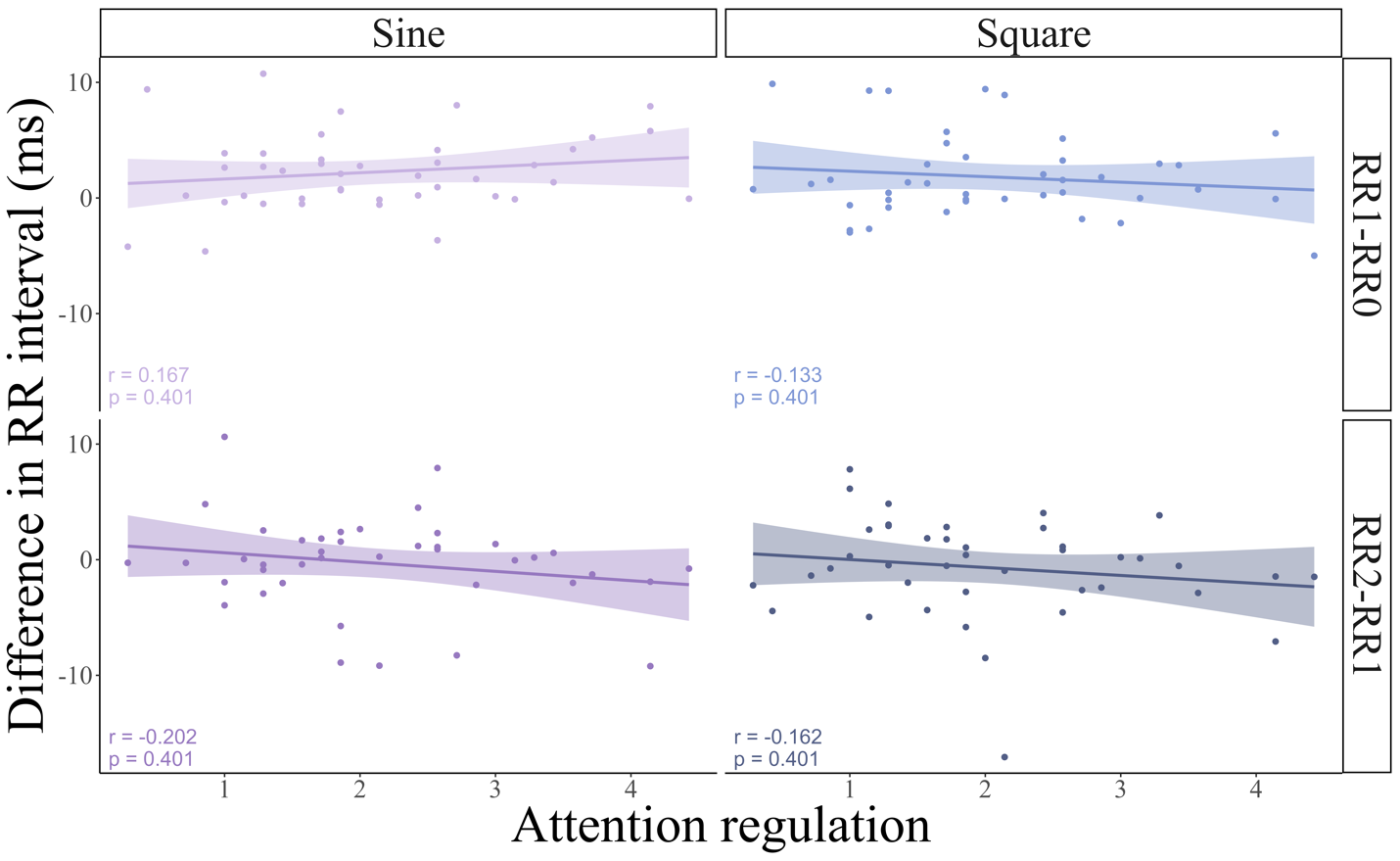

**Fig. A.7.**

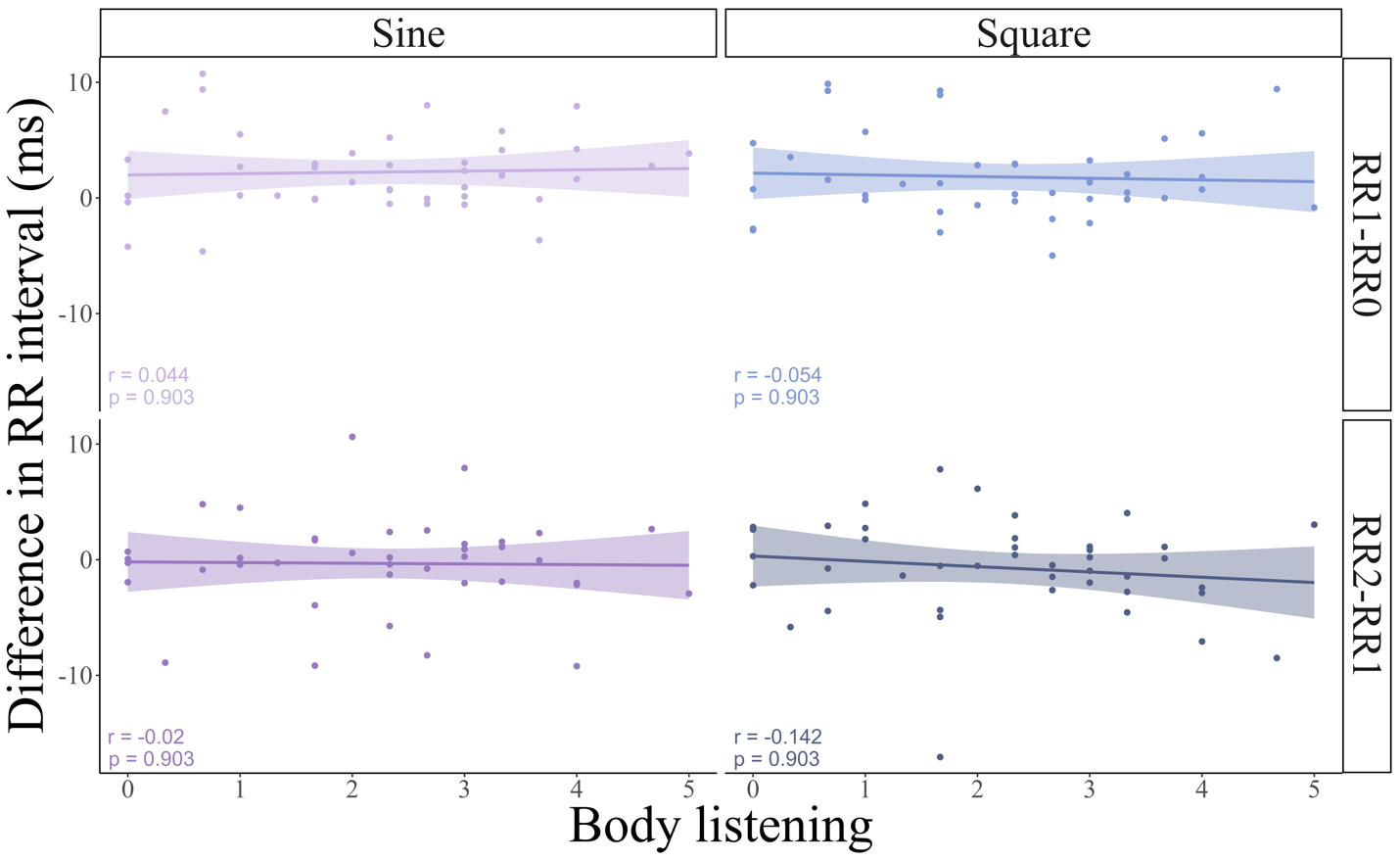

**Fig. A.8.**

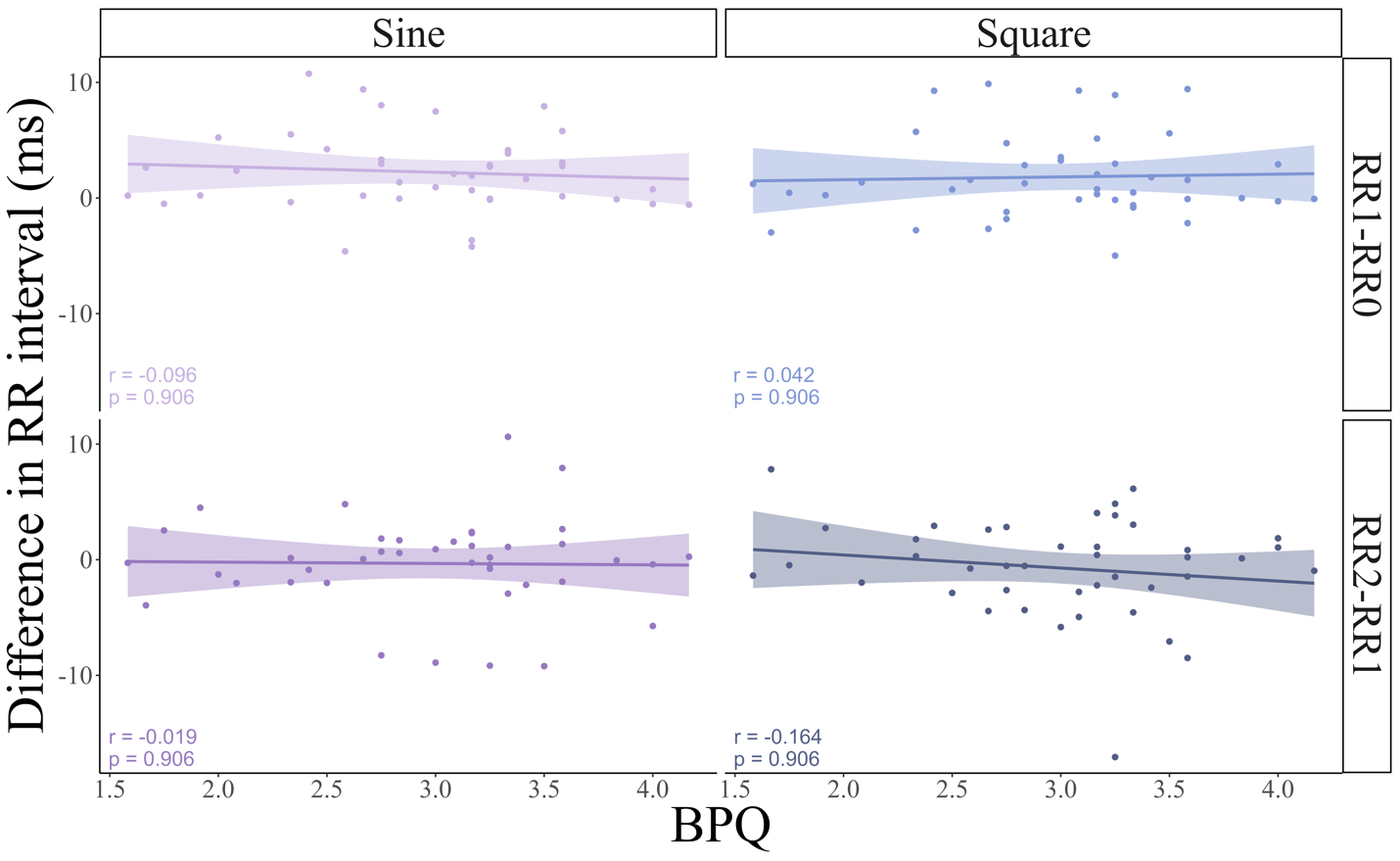

**Fig. A.9.**

**Table A.1.** Summary of estimated parameters in the model including RR1-RR0.

| Parameters | Estimate | | Est.Error | | l-95% CI | u-95% CI | Rhat | Bulk_ESS | Tail_ESS |
| --- | --- | --- | --- | --- | --- | --- | --- | --- | --- |
| Intercept | | 0.940 | | 0.714 | -0.483 | 2.347 | 1 | 3601.088 | 2710.158 |
| Sine | | -0.260 | | 0.381 | -1.026 | 0.496 | 1 | 3212.951 | 2947.383 |
| Square | | -0.896 | | 0.357 | -1.598 | -0.185 | 1 | 4243.063 | 3152.411 |
| RR1-RR0 | | -0.057 | | 0.089 | -0.235 | 0.114 | 1 | 2458.028 | 2378.784 |
| ECG_amplitude | | -0.073 | | 0.025 | -0.122 | -0.023 | 1 | 2786.141 | 2778.432 |
| Sex | | 0.621 | | 0.395 | -0.188 | 1.396 | 1 | 3094.149 | 2853.212 |
| Threshold | | -0.242 | | 0.290 | -0.792 | 0.330 | 1 | 4161.822 | 2972.766 |
| RR1-RR0*sine | | -0.012 | | 0.102 | -0.218 | 0.184 | 1 | 3028.029 | 2755.837 |
| RR1-RR0*square | | 0.035 | | 0.084 | -0.128 | 0.201 | 1 | 3115.126 | 3217.425 |
| RR1-RR0*HCT | | 0.052 | | 0.138 | -0.213 | 0.328 | 1 | 3003.385 | 2692.871 |

Notes:

R̂ = 1 for all parameters, the model has converged

Estimate: the estimated coefficient of each parameter

Est. Error: the standard error in the estimated coefficient of each parameter

l-95% CI: the lower limit of the 95% credible interval (CI)

u-95% CI: the upper limit of the 95% CI

R̂: indicators for determining convergence of Markov Chain Monte Carlo (MCMC) chain

Usually, the model is considered to have converged at R̂ ≤ 1.1

Bulk_ESS: the effective sample size calculated by normal rank

Tail_ESS: the effective sample size for the 5% and 95% quantile points

Intercept: Reference level of the objected variable excluding the effects of other variables

Sine: the difference between base and sine conditions for main effects of stimulus type

Square: the difference between base and square conditions for main effects of stimulus type

RR1-RR0: the main effect of RR1-RR0

ECG_amplitude: the main effect of the amplitude of ECG

Sex: the main effect of the subjects’ biological sex

Threshold: the main effect of the subjects’ threshold of the electrical stimuli

Score: the main effect of the participants’ scores of the HCT

RR1-RR0*sine: the difference between the base and sine conditions for the interaction between RR1-RR0 and stimulus type

RR1-RR0*square: the difference between the base and square conditions for the interaction between RR1-RR0 and stimulus type

RR1-RR0*HCT: the interaction between RR1-RR0 and the score of the HCT
